# Histone deacetylase inhibition expands cellular proteostasis repertoires to enhance neuronal stress resilience

**DOI:** 10.1101/2024.08.21.608176

**Authors:** Caitlin A. Taylor, Maya Maor-Nof, Wendy A. Herbst, Shihong Max Gao, Xinyue Li, Eyal Metzl-Raz, Aaron Hidalgo, Faria Zafar, Callista Yee, Birgitt Schüle, Meng C. Wang, Aaron D. Gitler, Kang Shen

**Affiliations:** Department of Biology, Stanford University, Stanford, CA 94305, USA; Department of Biology, Technion - Israel Institute of Technology, Haifa, Israel; Department of Genetics, Stanford University School of Medicine, Stanford, CA 94305, USA; Janelia Research Campus, Howard Hughes Medical Institute, Ashburn, VA, USA; Department of Pathology, Stanford University School of Medicine, Stanford, CA, 94305, USA; Howard Hughes Medical Institute, Stanford University, Stanford, CA 94305, USA; The Phil and Penny Knight Initiative for Brain Resilience, Stanford University, Stanford, CA, 94305 USA; Chan Zuckerberg Biohub - San Francisco, San Francisco, CA, 94158, USA

## Abstract

Neurons are long-lived, terminally differentiated cells with limited regenerative capacity. The maintenance of proteostasis and membrane protein trafficking is essential for neuronal function throughout development and aging. Cellular stressors, including endoplasmic reticulum (ER) protein folding stress and endosomal membrane trafficking stress, accumulate as neurons age and correlate with age-dependent neurodegeneration. However, how neurons coordinate stress responses across diverse cellular perturbations remains poorly understood. Here we demonstrate that histone deacetylases (HDACs) constrain the flexibility of neurons to engage distinct molecular pathways in response to different types of stress. Genetic or pharmacological inhibition of class I HDACs enhances neuronal resilience to both ER protein folding stress and endosomal membrane trafficking stress in *C. elegans* and mammalian neurons. RNA sequencing analyses in *C. elegans*, mouse, and human iPSC-derived neurons reveal that HDAC inhibition makes neurons more developmentally plastic and induces a permissive transcriptional state that likely represents partial cell fate reprogramming. These transcriptomic changes enable neurons to activate latent proteostasis pathways tailored to the specific nature of the stress. Given the growing excitement around partial reprogramming as a strategy to reset the epigenetic clock, our findings establish a direct link between epigenetic modulation and the neuronal stress response, pointing to new therapeutic strategies for enhancing neuronal resilience in aging and disease.

## Main

Cells achieve proteostasis through the regulation of protein synthesis, folding and degradation. Dysregulation of proteostasis is a hallmark of aging and contributes to neurodegenerative diseases (Kurtishi et al., 2019). Many redundant, interconnected molecular pathways have evolved to maintain robust proteostasis (Balch et al., 2008). The unfolded protein response (UPR) maintains homeostasis against endoplasmic reticulum (ER) stress and contains three parallel signaling pathways, IRE1, PERK, and ATF6. UPR pathways detect unfolded proteins and resolve ER stress by inducing the production of chaperones to improve protein folding and by decreasing protein folding load (Wiseman et al., 2022). Acting in parallel to the UPR, ER-associated degradation (ERAD), promotes the clearance of misfolded protein in the ER (Wiseman et al., 2022). Similarly, extensive redundancy exists in the endo-membrane system that controls protein trafficking, distribution, recycling, and degradation (Lippincott-Schwartz et al., 2000). It remains unexplored how multiple proteostasis and membrane trafficking pathways are utilized during aging or neurodegeneration, or whether manipulating these pathways improves neuronal resilience to stress and achieves rejuvenation. Chromatin modification by histone deacetylase complexes (HDACs) is a powerful mechanism that regulates the cellular transcriptome during development (Seto & Yoshida, 2014). Interestingly, pharmacological or genetic inhibition of HDACs has shown neuroprotective effects at cellular and behavioral levels (Guan et al., 2009; Choong et al., 2016; Suelves et al., 2017; Janczura et al., 2018). Despite these compelling results, the underlying molecular mechanisms are not fully understood. Here we probe how inhibition of class I HDACs promotes rejuvenation across biological contexts and identify a set of resilience pathways that enable neurons to deal with diverse stressors.

### Loss of SIN-3/HDAC rescues neurons from endosomal and ER stress

To explore the molecular pathways utilized by neurons to cope with cellular stress, we focused on the *C. elegans* sensory neuron PVD. PVD displays an elaborate dendritic arbor (Figure 1a) that requires proper trafficking of membrane proteins through the secretory and endosomal systems (Figure 1b-c,j) (Heiman & Bülow, 2024). The loss of RAB-10 recycling endosomes results in a dramatic reduction of dendritic branches (Figure 1b,e) (Taylor et al., 2015; Zou et al., 2015), and we harnessed this phenotype to identify factors that improve neuronal resistance to membrane trafficking stress. We performed an unbiased forward genetic suppressor screen and identified two alleles of *sin-*3 that increased dendritic branch number in the *rab-10* null mutant background (Figure S1a-c). Two additional *sin-3* alleles, including a full deletion of the coding region by CRISPR-Cas9, also increased branch number in *rab-10* mutants (Figure S1a-c). Cell-specific deletion of *sin-3* in the PVD precursor cell suppressed the *rab-10* phenotype (Figure S1d-e), while overexpression of SIN-3 in all tissues or neurons abolished this suppression (Figure S1f-g), demonstrating that SIN-3 acts cell-autonomously. Thus, loss of SIN-3 is sufficient to partially restore the elaborate PVD neuron dendritic arbor in the absence of RAB-10.

**Figure 1.**
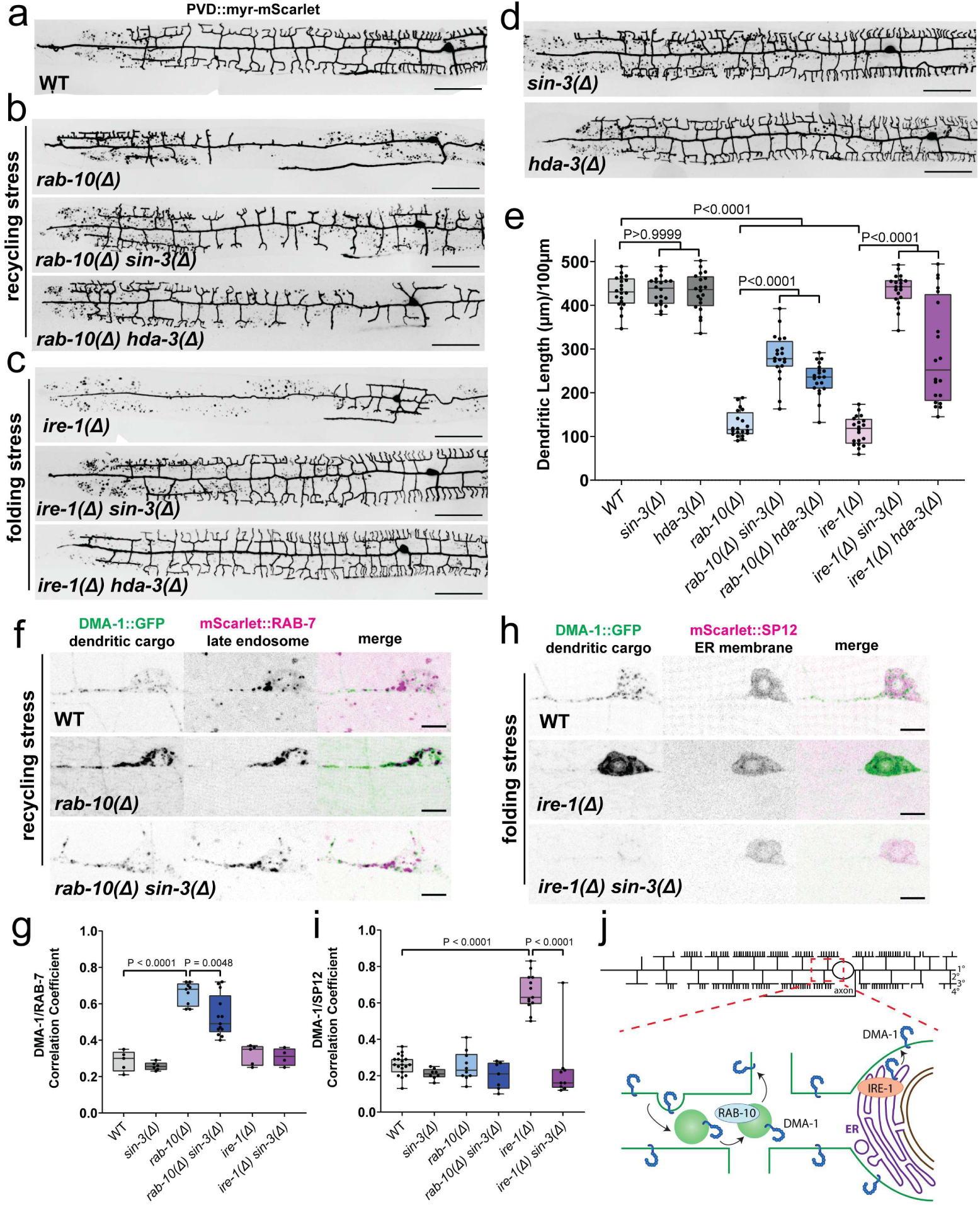
Loss of *sin-3/hda-3* protects against *rab-10* recycling stress and *ire-1* folding stress. (a) Representative image of wild-type PVD morphology. (b) Representative images of PVD morphology in *rab-10* single mutants and double mutants of *rab-10* with *sin-3* and *hda-3*. (c) Representative images of PVD morphology in *ire-1* single mutants and double mutants of *ire-1* with *sin-3* and *hda-3*. (d) Representative images of PVD morphology in *sin-3* and *hda-3* single mutants. Scale bars in (a-d) are 50 µm. (e) Quantification of total dendritic length per 100 µm of primary dendrites for genotypes shown in a-d. n=20 for all genotypes. (f) Representative images of endogenous DMA-1::GFP and endogenous mScarlet::RAB-7 in PVD somatodendrite region in wild-type (n=17), *rab-10* single mutants (n=9), and *rab-10 sin-3* double mutants (n=13). Scale bars are 5 µm. (g) Quantification of Pearson’s correlation coefficient, indicating degree of co-localization between DMA-1::GFP and mScarlet::RAB-7 for genotypes shown in f and additional genotypes. (h) Representative images of DMA-1::GFP and endogenous mScarlet::SP12 in PVD somatodendrite region in wild-type (n=19), *ire-1* single mutants (n=16), and *ire-1 sin-3* double mutants (n=9). Scale bars are 5 µm. (i) Quantification of Pearson’s correlation coefficient, indicating degree of co-localization between DMA-1::GFP and mScarlet::SP12 for genotypes shown in h and additional genotypes. (j) Schematic of a wild-type PVD dendrite (top) and routes of DMA-1 trafficking in the soma and primary dendrite (bottom). All statistical comparisons were performed using one-way ANOVA with Tukey’s correction for multiple comparisons.

Because of the striking ability of *sin-3* to suppress the *rab-10* dendrite loss phenotype, we next asked if *sin-3* mutations could suppress a neuronal phenotype caused by a different cellular stress. All eukaryotic cells monitor proper protein folding in the endoplasmic reticulum (ER) and respond to ER stress by activating a network of signaling pathways called the unfolded protein response (UPR), and hallmarks of ER stress have been reported in many diseases (Hoozemans et al., 2009; Nijholt et al., 2012; Hetz & Saxena, 2017; Singh et al., 2024). Inositol-requiring enzyme (IRE1) is an ER stress sensor essential for the UPR in yeast, plants, and animals. *C. elegans ire-1* mutations severely disrupt the PVD dendritic arbor (Figure 1c,e) (Wei et al., 2015; Salzberg et al., 2017). Remarkably, *sin-3* mutation was able to completely rescue the dendritic arbor in *ire-1* deletion mutants (Figure 1c,e). Thus, *sin-3* deficiency potently reversed the severe dendritic branching phenotype elicited by two distinct perturbations (trafficking stress and ER stress). We did not observe PVD phenotypes in *sin-3* single mutant animals (Figure 1d-e), suggesting that SIN-3’s function is dispensable during normal development but becomes important upon cellular perturbations such as *rab-10* or *ire-1* deficiency.

SIN-3 is a highly conserved scaffolding and transcriptional corepressor protein that interacts with histone deacetylases (HDACs). SIN-3/HDAC complexes repress gene expression by deacetylating nucleosomes and altering chromatin structure (Seto & Yoshida, 2014). We first asked if SIN-3’s effects on PVD dendritic growth depended on HDACs. We tested mutations in six different *C. elegans* HDAC genes and found that loss of only the class I HDAC, *hda-3*, was able to suppress the branching defects of the *ire-1* mutant (Figure 1c,e; Figure S1h-i,), suggesting that SIN-3 and HDA-3 function together. We identified two additional conserved components of the SIN-3 complex (ARID-1/ARID4 and ATHP-1/PHF12) whose mutants were also able to suppress the branching defects of *ire-1* mutants (Figure S1h-j), further supporting the notion that SIN-3 and HDA-3 function as a transcriptional repressor complex.

Most HDAC family members regulate transcription through deacetylating histones, but a few have been shown to work on cytoplasmic proteins such as tubulins (Hubbert et al., 2002; Rivieccio et al., 2009). We examined subcellular localization by generating GFP knockins at the endogenous loci with CRISPR-Cas9. SIN-3 and HDA-3 were both constitutively expressed and nuclear-restricted in PVD neurons throughout development, indicating that they likely regulate chromatin (Figure S1k-l). SIN-3 and HDA-3 localization was unperturbed in *rab-10* and *ire-1* mutants (Figure S1k-l), indicating stress itself does not alter localization of the SIN-3/HDA-3 complex. Together, these results suggest that a SIN-3/HDA-3 complex acts in the nucleus (Figure S1j) to limit the ability of neurons to cope with membrane trafficking or ER stress.

Because *sin-3* deficiency rescued neuronal phenotypes elicited by strong loss-of-function alleles of *rab-10* or *ire-1*, it follows that *sin-3* must activate alternative cellular pathways for PVD dendrite branching. We hypothesized that loss of SIN-3 activated a membrane trafficking “backup pathway” that is normally quiescent and is only implemented under stressed conditions. One possibility is that such a “backup pathway” involves a cell-fate change. However, loss of the transcription factors *mec-3* or *unc-86*, both of which are required for PVD specification (Way & Chalfie, 1989; Tsalik et al., 2003; Smith et al., 2010), entirely abolished dendritic arbors in the *sin-3* mutant background (Figure S1m), indicating that *sin-3* mutations did not change PVD’s terminal identity. We also found that the loss of *sin-3* did not alter the guidance receptors used for PVD growth and patterning. DMA-1 is a transmembrane receptor that promotes dendrite branching and growth in PVD neurons (Liu & Shen, 2011). Deleting *dma-1* abolished dendrites in *sin-3* mutants, suggesting that *sin-3* mutants did not completely alter the molecular program for branching (Figure S1n).

How could a potential backup pathway resolve two different types of cellular stress and promote dendrite growth? To address this, we next analyzed how *sin-3* rescued the *rab-10* and *ire-1* mutant phenotypes by examining the intracellular trafficking of the key dendrite promoting molecule DMA-1 (Eichel et al., 2022), which is normally found in RAB-10-positive vesicles (Supplemental Movie 1) (Shi et al., 2024). In *rab-10* mutants, DMA-1 was largely missing from the dendrite and abnormally accumulated in the soma and proximal primary dendrite in RAB-7-positive compartments (Figure 1f-g). In the *rab-10 sin-3* double mutant, mistargeting of DMA-1 was significantly resolved (Figure 1f-g and Supplemental Movie 2). Additionally, the excessive accumulation of RAB-7 endosomes was reduced, suggesting that *sin-3* likely affects trafficking to RAB-7 endosomes rather than specifically regulating DMA-1 (Figure 1f).

DMA-1 localization was also defective in *ire-1* mutants. However, unlike in *rab-10* mutants, the *ire-1* mutant showed drastically reduced DMA-1 in the dendrite and a strong accumulation in the somatic compartment (Figure 1h-i). Consistent with IRE-1’s function in the UPR, somatic DMA-1 colocalized with the ER marker SP12, suggesting that DMA-1 accumulates in the ER in *ire-1* mutants (Figure 1h-i). In *ire-1 sin-3* double mutants, this ER-accumulation of DMA-1 was completely resolved (Figure 1h-i). While *rab-10* and *ire-1* PVD neurons both exhibit dendritic branching defects, the underlying cell biological phenotypes differ, and remarkably *sin-3* deficiency rescued both phenotypes. Together, these data provide evidence that PVD neurons have the capacity to engage alternative protein-processing pathways to resolve diverse membrane trafficking and ER deficits, but perplexingly these compensatory pathways can only be utilized in the absence of *sin-3*.

### Protective effects of HDAC inhibition in ER stress rely on multiple proteostasis pathways

After establishing that the loss of *sin-3/hda-3* provided protection against multiple developmental stressors, we next sought to understand the molecular mechanisms of this protective effect. Previous studies across species have shown that HDACs mediate thousands of transcriptional changes, which presents a unique challenge when trying to understand the specific mechanisms that underlie the benefits of HDAC inhibition. We were able to leverage several advantages of the *C. elegans* PVD system to address this challenge. Our direct observation of relevant dendritic cargoes (DMA-1) suggested that the loss of *sin-3* largely resolved a key readout of ER stress in *ire-1* mutants. DMA-1 accumulated in the ER of *ire-1* single mutants, and this accumulation was cleared in *ire-1 sin-3* double mutants. This led us to hypothesize that the beneficial effect of *sin-3* deficiency likely relied upon cellular pathways related to protein folding, protein clearance, and ER function. We curated a list of genes involved in these pathways and performed a candidate screen to identify genes required for the protective effect of *sin-3* deficiency (Figure S2a). We generated triple mutants of *ire-1, sin-3,* and loss-of-function alleles of candidate genes and looked for mutants that abolished the protective effect of *sin-3*. We identified seven relevant genes which we refer to as “backup pathway” genes (Figure S2b). In triple mutants of each backup pathway gene with *ire-1 sin-3*, dendritic branch growth was dramatically reduced (Figure 2a-b, Figure S2b-c). These genes were involved in the unfolded protein response/UPR (*atf-6*/Atf6 and *pek-1*/Eif2ak3), integrated stress response/ISR (*eif-2α*/Eif2s1), ER-associated degradation/ERAD (*sel-1*/Sel1l and *sel-11*/Syvn1), protein folding (*hsp-3*/Hspa5), and a chaperone-interacting neuroprotective factor (*manf-1*/Manf), and all of these genes functioned cell-autonomously (Figure S2c-e). Many other proteostasis genes tested in the candidate screen were not required for the protective effect, including additional ERAD ubiquitin ligases (*hrdl-1, marc-6,* or *rnf-5*), UPR transcription factors (*xbp-1*), and chaperone proteins (*hsp-4)* (Figure S2b). Together these results suggest that specific proteostasis pathways are required in PVD for the protective effect of *sin-3* deficiency.

**Figure 2.**
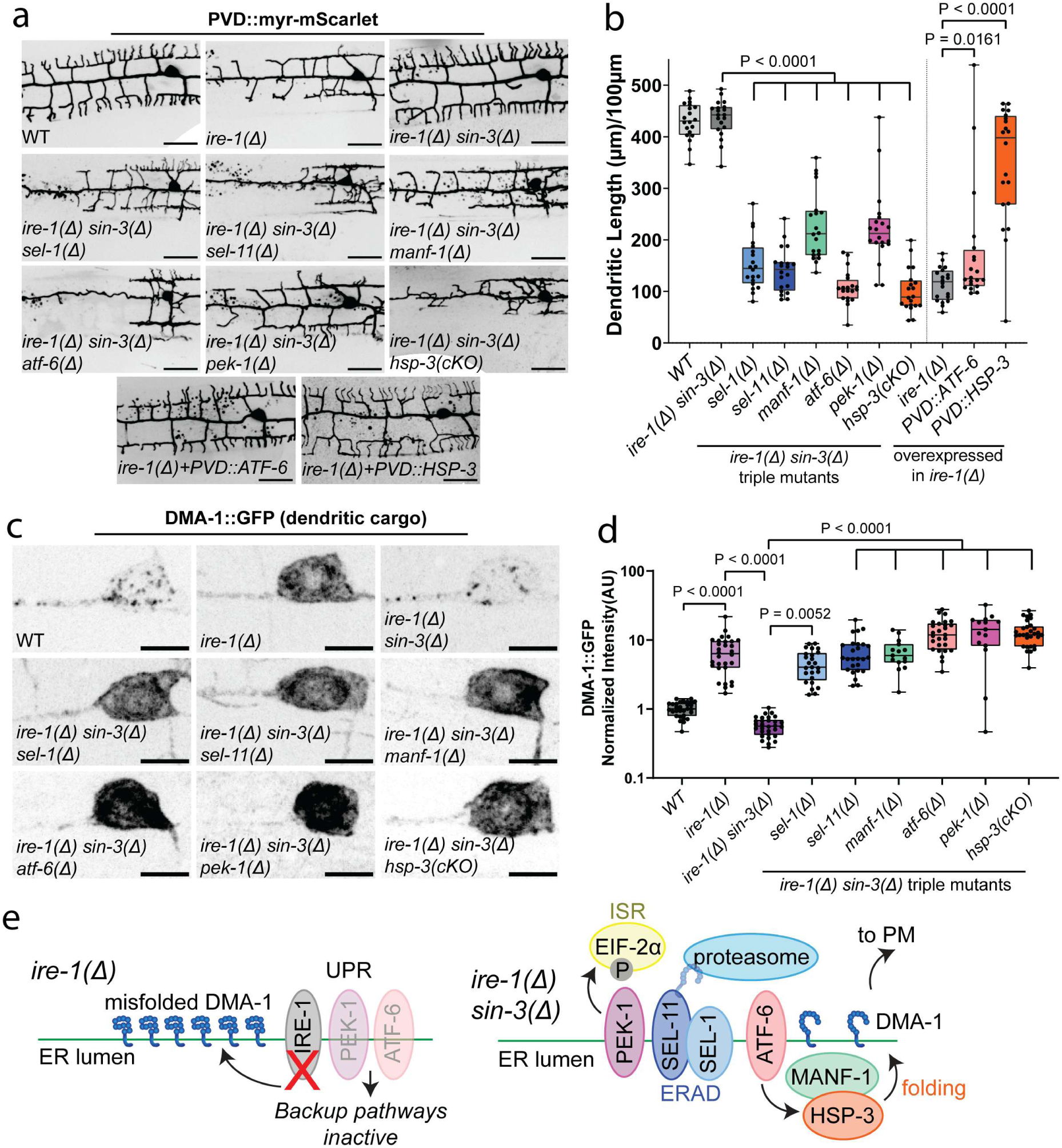
Multiple proteostasis pathways are required for the protective effect of *sin-3*. (a) Representative images of PVD morphology in wild-type, *ire-1* single mutant, *ire-1 sin-3* double mutant, triple mutants of *ire-1 sin-3* with *sel-1, sel-11, manf-1, atf-6, pek-1,* or *hsp-3,* and single mutants of *ire-1* with PVD-specific overexpression of ATF-6 or HSP-3. Scale bars are 25 µm. (b) Quantification of total dendritic length per 100 µm of primary dendrites for genotypes shown in a. n=20 for all genotypes. (c) Representative images of endogenous DMA-1::GFP in soma of wild-type (n=30), *ire-1* single mutant (n=30), *ire-1 sin-3* double mutant (n=30), and triple mutants of *ire-1 sin-3* with *sel-1* (n=25)*, sel-11* (n=26)*, manf-1* (n=15)*, atf-6* (n=27)*, pek-1* (n=15), or *hsp-3* (n=30). Scale bars are 5 µm. (d) Quantification of DMA-1::GFP intensity. All genotypes were normalized to the median of wild-type and data are plotted on a log-scale. (e) Schematic of trafficking pathways in *ire-1* mutant leading to misfolded protein (left) and backup pathways activated in *ire-1 sin-3* double mutant (right). All statistical comparisons were performed using one-way ANOVA with Tukey’s correction for multiple comparisons.

Importantly, none of the backup pathway genes were required for normal development: single mutants did not generate dendritic phenotypes, and a sextuple mutant of *manf-1 sel-1 sel-11 atf-6 pek-1 hsp-3* also showed normal dendritic development (Figure S2f-g). Similarly, none of these genes were required for DMA-1 localization in an unstressed background; single mutants showed no change in DMA-1 localization (Figure S2h-i). These data suggest that these genes represent a true “backup pathway”: none of these pathways are required during normal development. Further, they cannot be recruited to support branch growth and resolve ER stress in the absence of *ire-1* unless *sin-3* activity is inhibited.

To define how upregulating these alternative pathways via HDAC inhibition helped restore the PVD neuron dendritic arbor in the absence of *ire-1*, we examined the localization of DMA-1::GFP. In triple mutants of *ire-1 sin-3* with backup pathway genes involved in ERAD (*sel-1*, *sel-11)*, folding (*hsp-3, manf-1*), and UPR (*atf-6, pek-1*), we observed a dramatic accumulation of DMA-1 in the cell body (Figure 2c-d), which likely represents misfolded, ER-trapped protein that cannot be degraded. In the *sin-3* mutant, three pathways are utilized to improve DMA-1 proteostasis: clearance of misfolded protein relies on ERAD (*sel-11* and *sel-1)*, improved folding is mediated by UPR and chaperone proteins (*atf-6, hsp-3*, and *manf-1*), and translational load is reduced by activation of the ISR (*pek-1* and *eif-2α*) (Fig 2e). We next sought to directly measure ERAD activity using a well-established ERAD substrate (Finger, et al 1993). We expressed CPL-1[W32A,Y35A]::mScarlet in PVD (Figure S3a). As expected, ERAD mutants *sel-1* or *sel-11* showed increased signal (Fig S3b-c). Interestingly, we observed a strong reduction of CPL-1[W32A,Y35A] signal in *sin-3* single mutants (Figure S3b-c), which suggested that ERAD activity was higher in the *sin-3* mutants. To confirm the involvement of ERAD, we constructed double mutants of *sin-3 sel-1* or *sin-3 sel-11* and found that CPL-1[W32A,Y35A] levels were restored to a higher level (Figure S3b-c), indicating that lower CPL-1[W32A,Y35A] levels in *sin-3* mutants were a result of increased clearance by ERAD.

We next tested whether the backup pathway genes required for dendritic growth in *ire-1 sin-3* were individually sufficient to rescue *ire-1* dendritic phenotypes. Overexpression of the chaperone protein HSP-3 rescued the dendritic growth phenotype of *ire-1* mutants, while overexpression of the UPR regulator ATF-6 showed a weaker but significant rescue (Figure 2a-b). Overexpression of SEL-11, SEL-1, PEK-1, or MANF-1 yielded no increase in dendritic growth in *ire-1* single mutants (Figure S2j-k). While our loss-of-function data indicate that the beneficial effect of *sin-3* mutation requires the coordinated regulation of multiple arms of the backup pathway, these overexpression data suggest that upregulation of HSP-3, and to a lesser extent ATF-6, is individually capable of protecting neurons from ER stress.

We next asked how the backup pathway genes that were identified in *C. elegans* were transcriptionally regulated by HDACi. We generated transcriptional reporters for *manf-1*, *hsp-3*, *sel-11*, and *atf-6* by knocking in a P2A-NLS-mNeonGreen cassette at each endogenous locus. We observed two different transcriptional patterns with these reporters. All four genes were strongly induced by the loss of *ire-1* (Figure S3d-h), suggesting that ER stress alone drives increased expression of backup pathway genes. Two of the genes (*hsp-3* and *manf-1*) were maintained at high levels in *ire-1 sin-3* double mutants (Figure S3d,e-f), while two (*sel-11* and *atf-6*) were attenuated in the *ire-1 sin-3* double mutants (Figure S3d,g-h). We hypothesize that the attenuated genes track ER stress status (elevated only in the presence of misfolded protein), while the two genes that remain high represent a sustained induction of backup pathways even after ER stress has been apparently resolved. These data indicate that UPR and ERAD were activated to some extent in the *ire-1* single mutant but were insufficient to resolve the ER stress. We hypothesize that the loss of *sin-3* altered the transcriptional dynamics of backup pathway genes, allowing for the prolonged expression of protective backup pathway genes.

### Activation of alternate endosomes alleviates endosomal recycling stress

Our data suggest that a network of genes involved in protein folding and degradation, including ERAD genes, Manf, BiP/Hspa5, and the PERK and Atf6 arms of the UPR, are required to protect neurons from ER stress. But are these same genes also required to protect neurons from endolysosomal stress, which has different underlying cell biological defects? We constructed triple mutants of *rab-10, sin-3* and the ER stress related backup pathway genes. Surprisingly, loss of these ER stress backup pathway genes had little effect on the *rab-10, sin-3* double mutant (Figure S4a-b). Loss of *sel-1, sel-11, manf-1,* or *pek-1* did not affect branch number in the *rab-10 sin-3* double mutant, while loss of *atf-6* caused a minor reduction in branches but did not completely abolish the protective effect as it did with *ire-1.* Together, these results suggest that HDAC inhibition rescues the *rab-10* PVD neuron branching phenotype by activating a different cellular response.

We sought to identify this alternative protective response and to define the cellular and molecular mechanisms by which *sin-3* deletion promotes growth in *rab-10* mutant neurons. We labeled three different populations of endosomes to monitor membrane trafficking in *rab-10* and *sin-3* mutants. We visualized early endosomes (RAB-5), late endosomes (RAB-7), and an additional species of recycling endosomes (RAB-11.1) in PVD neurons using cell-specific, endogenous GFP knockins (Figure S5a-e). In *rab-10* mutants, we observed a depletion of RAB-5 early endosomes (Figure S5a-b) and an accumulation of both RAB-7 late endosomes and RAB-11.1 endosomes (Figure S5a,c-d). We hypothesize that this altered distribution represents an “endolysosomal traffic jam” – in the absence of the preferred recycling endosome RAB-10, material was mis-sorted. This resulted in a cascade of dysregulation throughout the interconnected endosomal system, leading to a depletion of RAB-5 early endosomes and an increase in both RAB-7 late endosomes (Supplemental Movie 2) and RAB-11.1 recycling endosomes (Supplemental Movie 3).

Both the depletion of RAB-5 early endosomes and the accumulation of RAB-7 late endosomes were rescued in *rab-10 sin-3* double mutants, suggesting that *sin-3* deficiency helped to clear the accumulation of mis-sorted cargos and restore the balance between early and late endosomes (Figure S5a,c). In contrast, the accumulation of RAB-11.1 recycling endosomes was unaffected in *rab-10 sin-3* double mutants (Figure S5a,d). The sustained higher level of RAB-11.1 suggested two possibilities. First, RAB-11.1 accumulation could represent a detrimental consequence of the *rab-10* mutation that the loss of *sin-3* was unable to resolve. Alternatively, the increased level of RAB-11.1 could represent a compensatory response that was beneficial for branch growth in the *sin-3* mutant.

To distinguish these possibilities, we inactivated RAB-11.1 by expressing a dominant-negative form of RAB-11.1[S25N] in PVD neurons. Inhibiting RAB-11.1 caused a severe loss of dendritic branches in the *rab-10 sin-3* double mutant (Figure 3a-b), indicating that RAB-11.1 endosomes are required for the rescuing effect of *sin-3* deficiency.

**Figure 3.**
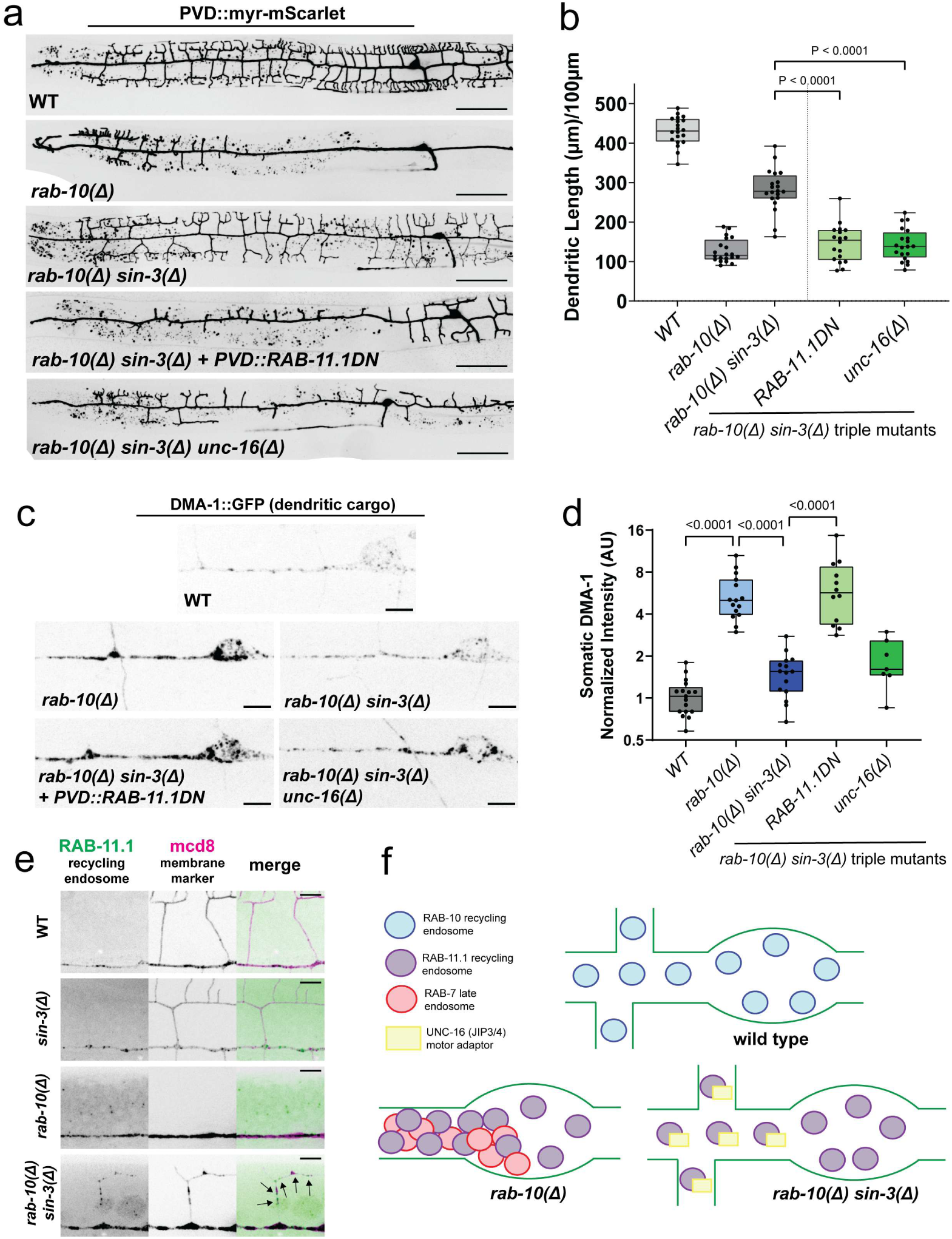
Alternative endosomal trafficking is required for protective effect of *sin-3*. (a) Representative images of PVD morphology in wild-type, *rab-10* single mutant, *rab-10 sin-3* double mutant, and triple mutants of *rab-10 sin-3* with RAB-11.1(DN) or *unc-16*. Scale bars are 50 µm. (b) Quantification of total dendritic length per 100 µm of primary dendrites for genotypes shown in a. n=20 for all genotypes. (c) Representative images of endogenous DMA-1::GFP in soma of wild-type (n=17), *rab-10* single mutant (n=15), *rab-10 sin-3* double mutant (n=15), and triple mutants of *rab-10 sin-3* with RAB-11.1(DN) (n=12) or *unc-16* (n=7). Scale bars are 5 µm. (d) Quantification of DMA-1::GFP intensity. All genotypes were normalized to the median of wild-type and data are plotted on a log-scale. (e) Representative images of endogenous GFP::RAB-11.1 localization in higher order dendrites. (f) Schematic of endosomal trafficking in wild-type, *rab-10* single mutant, and *rab-10 sin-3* double mutant. All statistical comparisons were performed using one-way ANOVA with Tukey’s correction for multiple comparisons.

Inhibiting RAB-11.1 had no effect on dendritic branching in wild-type animals but fully abolished dendritic branches in the *rab-10* single mutant (Figure S4c-d,f). This suggests that RAB-11.1 functions in parallel to RAB-10 but is unable to support full branch growth in the absence of *rab-10*. We also examined the subcellular localization of the dendritic guidance receptor DMA-1, which is mis-localized in *rab-10* single mutants. We found that in *rab-10, sin-3*, RAB-11.1(DN) triple mutants, DMA-1 once again accumulated in internal structures in the soma and primary dendrite (Figure 3c), resulting in an overall increase in DMA-1 intensity (Figure 3d). This suggests RAB-11.1 functions to recycle or clear DMA-1 in the *rab-10 sin-3* double mutant.

While the abundance of RAB-11.1 endosomes was increased in the *rab-10* single mutant, this increase in RAB-11.1 supported only a small number of anterior dendrites and was insufficient to generate a full dendritic arbor. By contrast, when *sin-3* was deleted in *rab-10* mutants, RAB-11.1’s role was potentiated, and it became capable of largely compensating for the loss of *rab-10*. How might this potentiation occur? While RAB-11.1 levels were not further altered by the loss of *sin-3,* we asked whether the subcellular localization and movements of RAB-11.1 endosomes were altered by the loss of *sin-3*. In wild-type animals, we rarely observed RAB-11.1 puncta in the higher-order dendrites (Figure 3e). Instead, RAB-11.1 showed processive movement along the primary dendrite (Supplementary Video 4). In striking contrast, in *rab-10, sin-3* double mutant animals, we observed both stable and dynamic RAB-11.1 puncta in most secondary dendrites (Figure 3e, Supplementary Video 4). This RAB-11.1 behavior recreated the normal dynamics of RAB-10 endosomes in wild-type animals (Supplementary Video 5). We hypothesized that this change in trafficking behavior could be mediated by motor adaptors such as the JNK-interacting proteins 3 and 4 (JIP3/JIP4), which mediate the association of cargos with the molecular motors kinesin and dynein (Fu and Holzbaur, 2014). To test the role of motor adaptors, we generated a triple mutant between *rab-10, sin-3*, and *unc-16*, the sole *C. elegans* homolog of JIP3/4. The loss of *unc-16* alone did not cause dendritic defects in otherwise wild-type animals (Figure S4c,f) but resulted in a strong reduction in dendritic branches in the *rab-10 sin-3* double mutant (Figure 3a-b), suggesting that *unc-16* is required for the protective effects of *sin-3* deficiency. We hypothesize that UNC-16 mediates the altered trafficking of RAB-11.1 in *sin-3* mutants, and future studies will explore the mechanisms by which *sin-3* regulates UNC-16 to mediate endosomal trafficking.

These data suggest that the loss of *sin-3* enables neurons to utilize a RAB-11.1/UNC-16 trafficking pathway (Figure 3f). We next asked whether this endosomal backup pathway was required to rescue ER stress phenotypes. Just as the ER stress backup pathway genes were not required for *sin-3* to rescue the *rab-10* phenotype, RAB-11.1 and UNC-16 were also dispensable for *sin-3* to rescue the *ire-1* phenotype – in *ire-1*, *sin-3*, and RAB-11.1(DN) or *unc-16* triple mutants, dendrites were similar to wild-type controls (Figure S4e-f). This suggests that distinct stressors, including ER stress or endosomal stress, require different compensatory responses that are both mediated by HDAC inhibition.

### HDAC inhibition protects against ER stress in mouse cortical neurons

Because of the potent beneficial effect of inhibiting the SIN-3/HDA-3 complex on dendritic arbor development in stressed *C. elegans* neurons, we next asked if HDAC inhibitors could protect against additional stressors in mammalian neurons. We used mouse cortical neurons as a model to examine neuronal and axonal degeneration in response to ER stress. We cultured E16.5 mouse primary cortical neurons for three days to allow process outgrowth and then treated with 1μg/mL tunicamycin (Figure 4a). Tunicamycin causes accumulation of unfolded glycoproteins in the ER, leading to ER stress (Oslowski & Urano, 2011). Treatment of mouse cortical neurons with tunicamycin caused numerous tubulin βIII puncta in the neurites, indicating that microtubules disassembled, and the axons underwent degeneration (Figure 4b,d).

**Figure 4.**
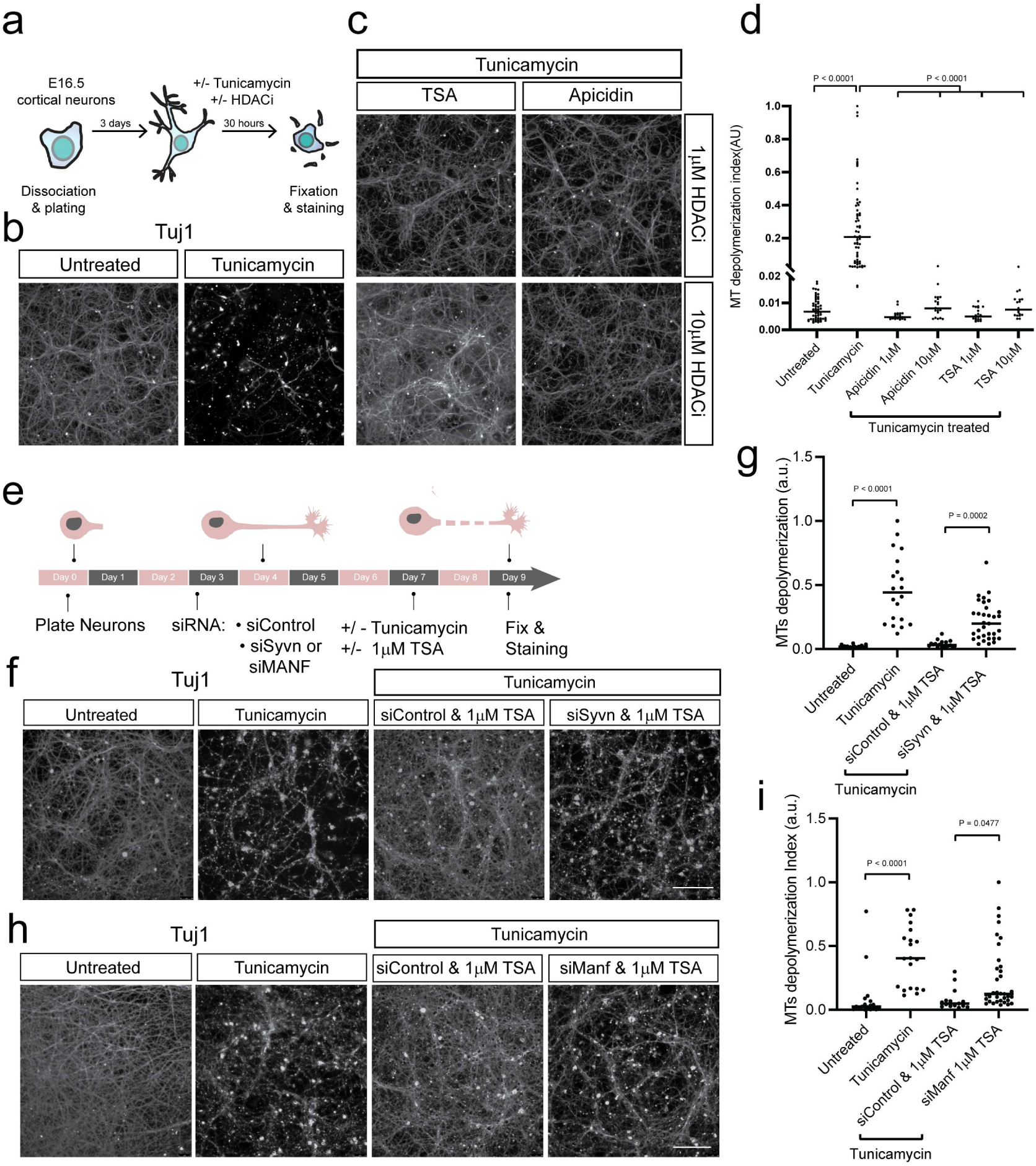
HDACi protects against tunicamycin-induced degeneration in mouse cortical neurons. (a) Schematic of tunicamycin and HDACi treatment of mouse cortical neurons. (b) Representative images of tubulin βΙΙΙ staining in mouse cortical neurons treated with or without tunicamycin. (c) Representative images of tubulin βΙΙΙ staining in mouse cortical neurons co-treated with tunicamycin and either apicidin or TSA. (d) Quantification of microtubule depolymerization index determined from tubulin βΙΙΙ staining. (e) Schematic of siRNA experiment in mouse cortical neurons. (f) Cortical primary neurons treated with both tunicamycin and TSA after Syvn1 or non-targeting siRNA were added. (g) Quantification of microtubule depolymerization index in Syvn1 knockdown, determined from tubulin βΙΙΙ. (h) Cortical primary neurons treated with both tunicamycin and TSA after Manf or non-targeting siRNA were added. (i) Quantification of microtubule depolymerization index in Manf knockdown, determined from tubulin βΙΙΙ staining. All scale bars are 100 µm. All statistical comparisons were performed using one-way ANOVA with Tukey’s correction for multiple comparisons.

To test whether HDAC inhibition could protect neurons from degeneration, we blocked HDAC activity with two HDAC inhibitors, apicidin and trichostatin A (TSA). TSA affects a wider range of HDAC isoforms, whereas apicidin targets specifically Class I HDACs, which include HDAC1, HDAC2, and HDAC3 (Bradner et al., 2010). We treated neurons with tunicamycin alone or co-treated with tunicamycin and 1μM and 10μM of TSA or apicidin for 30 hours and stained for tubulin βIII to assess neuronal survival and neurite integrity (Figure 4c-d). Tunicamycin treatment alone caused strong loss of neurites, but co-treatment with TSA or apicidin was sufficient to protect neurons from degeneration (Figure 4c-d) in a dose dependent manner (Figure S6a-c). The protective effect conferred by HDAC inhibition was evident for more than 36 hours following ER stress induction. Thus, HDAC inhibition enables neuronal resilience against the toxicity of tunicamycin-induced ER stress.

We next tested if ERAD and Manf were also required for HDAC inhibition’s protective effect against ER-stress induced degeneration in mouse neurons. We used small interfering RNA (siRNA) against Syvn1, the mammalian ortholog of *sel-11*, or Manf, the mammalian ortholog of *manf-1*. We treated primary cortical neurons with either non-targeting siRNA, Syvn1 siRNA, or Manf siRNA for 65 hours before the addition of tunicamycin and HDAC inhibitors, TSA and apicidin, for an additional 30h (Figure 4e). HDAC inhibition conferred protection against axonal degeneration and cell death in ER-stressed neurons treated with a control non-targeting siRNA (Figure 4f-i). In contrast, Syvn1 or Manf knockdown markedly attenuated the protective effect (Figure 4f-i). These results suggest that Syvn1 and Manf are important and highly conserved (from worms to mammals) components of resilience program to ER-stress.

Together, these data demonstrate that both ERAD and protein folding machinery are coordinately regulated as part of the protective pathway activated by HDAC inhibition in both *C. elegans* and mouse neurons. This conservation is perhaps surprising, as the two stress paradigms are quite different. In the *C. elegans* model, the ER stress is a developmental event affecting dendritic outgrowth, and the neurons do not experience cell death. In the mouse model, ER stress induces axonal degeneration and eventual cell death. The cell biological underpinnings of these events (dendritic development versus neuronal and axonal degeneration) are likely very different, but in both cases HDAC inhibition provides protection that depends on the same molecular players, Manf/*manf-1* and Syvn1/*sel-11*. While ER stress might elicit distinct consequences in different contexts, HDAC inhibition can provide protection in both dendrite development and neuronal degeneration.

### HDAC inhibition alters transcriptional state in *C. elegans*, mouse, and human neurons

HDACs are chromatin modifiers that play essential roles in the regulation of transcription in the physiological contexts of cell survival, growth, proliferation and differentiation (Seto & Yoshida, 2014). Characterizing the transcriptional targets of HDACs is essential to understanding the protective effects of HDAC inhibition. We identified molecular programs required for the protective effect of HDAC inhibition using hypothesis-driven reverse genetics in *C. elegans.* However, it was unclear whether these proteostasis genes were direct targets of HDACi. Additionally, given the broad effect of HDACi, it was unlikely that the effect of HDACi was specific to proteostasis pathways. We performed a series of RNA-sequencing experiments to understand the transcriptional regulation of backup pathway genes as well as to uncover additional transcriptional programs regulated by HDACi.

First, we examined how HDACi affected the transcriptomes of both stressed and unstressed mouse cortical neurons (Figure 5a). To determine the ideal time point after ER stress induction at which to assess the transcriptional programs involved in degeneration or protection, we added HDAC inhibitors at different time points after tunicamycin treatment and assessed the magnitude of axonal degeneration after 24 hours. We found that inhibiting HDACs within 7 hours of tunicamycin treatment provided robust protection against axonal degeneration. To define the transcriptional program induced by HDAC inhibition with or without ER stress, we extracted neurons at 7 hours following tunicamycin treatment in the presence or absence of HDAC inhibitors and performed RNA sequencing (RNA-seq) (Figure 5a).

**Figure 5.**
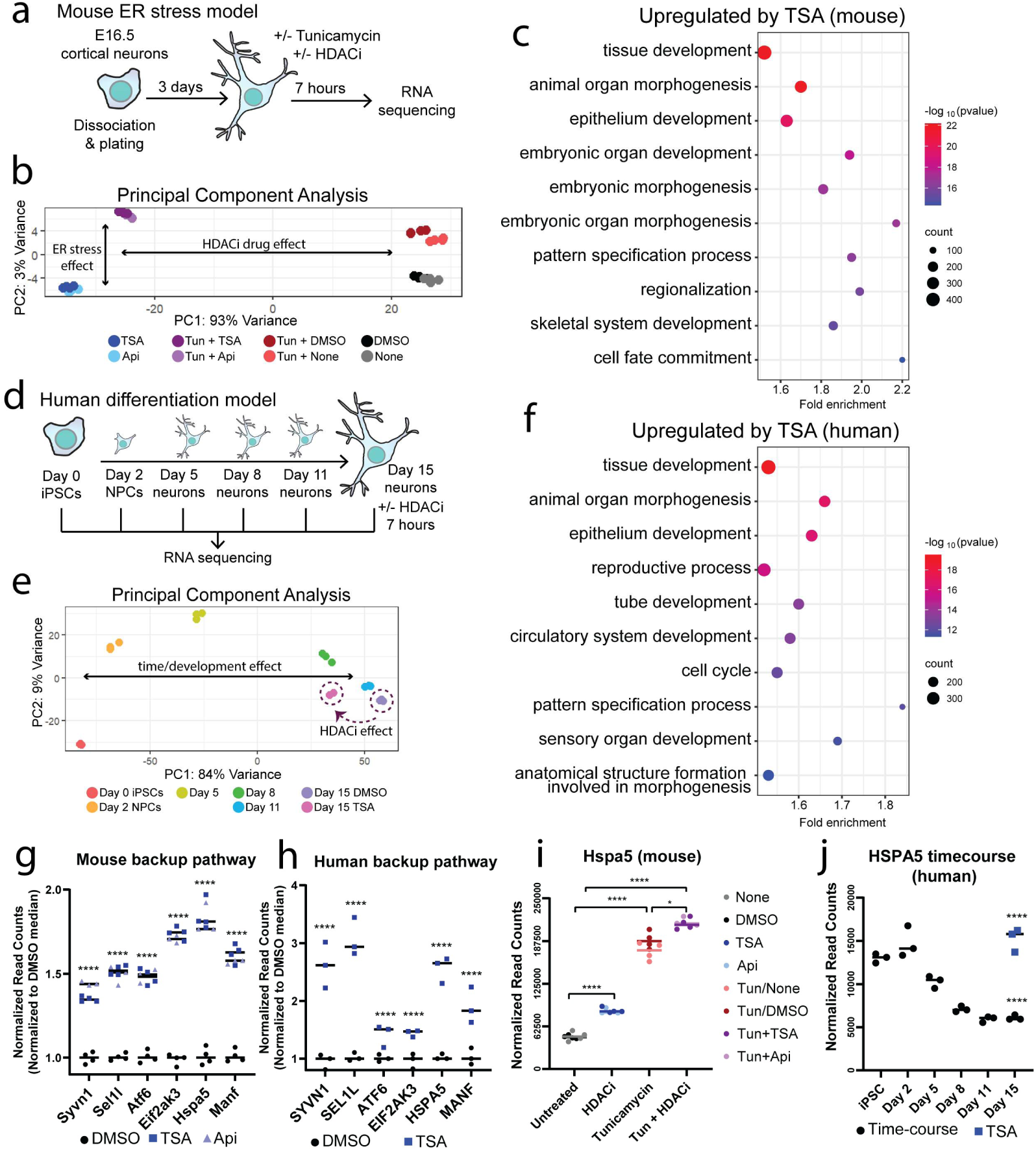
HDACi alters transcriptomic state in mouse and human neurons. (a) Schematic of RNA-sequencing experiment performed in mouse cortical neurons. (b) Principal component analysis of all drug treatment groups. (c) GO analysis of genes upregulated by TSA in mouse cortical neurons. (d) Schematic of RNA-sequencing experiment performed in human iPSC-derived neurons. (e) Principal component analysis of timecourse and HDACi treatment group. Day 15 conditions (with and without TSA) are circled. (f) GO analysis of genes upregulated by TSA in human iPSC-derived neurons. (g) Normalized read counts of all backup pathway gene expression in mouse cortical neurons treated with only DMSO, TSA, or apicidin. All replicates were normalized to the median of DMSO for each gene. **** represents p<0.0001. (h) Normalized read counts of all backup pathway gene expression in Day 15 human iPSC-derived neurons treated with DMSO or TSA. All replicates were normalized to the median of DMSO for each gene. **** represents p<0.0001. (i) Normalized read counts of Hspa5 expression in mouse cortical neurons treated with +/- tunicamycin and +/- HDACi. All conditions are plotted, while statistical comparisons shown are between DMSO and TSA groups. **** represents p<0.0001; * represents p = 0.0459. (j) Normalized read counts of HSPA5 expression across differentiation of human iPSC-derived neurons and in Day 15 neurons with or without TSA. Statistical comparisons shown are between Day 15 neurons with or without TSA (top) and between Day 15 neurons and Day 0 iPSCs (bottom). **** represents p<0.0001.

We used principal component analysis (PCA) to track the differential RNA profiles across treatment groups (Figure 5b), which revealed that the transcriptional changes were primarily explained by two principal components that reflected the changes following HDAC inhibition (PC1) and ER stress (PC2) (Figure 5b).

Rather than returning ER-stressed neurons to an “untreated” transcriptional state, the addition of HDACi pushed neurons into an orthogonal transcriptional state that was distinct from that of either untreated or ER-stressed neurons, suggesting an alternative cell state.

To define the nature of this orthogonal transcriptional state, we first analyzed the transcriptional program that was altered by HDACi without ER stress. Notably, we observed very few differentially expressed genes between neurons treated with the two different HDAC inhibitors (Figure S7A), suggesting that the transcriptional program regulated by either TSA or apicidin was highly consistent. We focused on the set of 6527 genes (3776 upregulated and 2751 downregulated) exhibiting the highest differential expression between TSA and DMSO treatment (Figure S7a). We performed Gene Ontology (GO) analysis on the differentially expressed genes following TSA treatment, which suggested a change in cellular identity. The most enriched pathways in the gene sets included “tissue development” and “animal organ morphogenesis,” demonstrating that HDACi activated many general developmental processes (Figure 5c). Interestingly, several GO terms involved in neuronal development were downregulated by TSA (Figure S7c). Taken together, these GO analyses suggest that TSA altered expression of neuronal and non-neuronal genes, resulting in a less strict cellular identity that we refer to as “partial reprogramming.” Interestingly, GO terms for proteostasis pathways were not enriched in the TSA treated neurons despite being essential for protection. However, the genes required for the protective effect in *C. elegans* were significantly upregulated by HDACi. The ERAD components Syvn1/*sel-11* and Sel1l/*sel-1*, UPR genes Atf6/*atf-6* and Eif2ak3/*pek-1*, and the chaperone-related genes Hspa5/*hsp-3* and Manf/*manf-1* were all significantly upregulated by both TSA and apicidin (Figure 5g). These data indicate that transcriptional regulation of the backup pathway genes was consistent with their function (i.e. required genes were also transcriptionally upregulated), but these pathways would not have been revealed by only focusing on the top regulated genes. Together our genetic and transcriptional analysis identified both specific pathways required for protection and indicated an overall state change toward “partial reprogramming,” which could contribute to the protective effect.

We next asked how HDACi affected the transcriptional programs induced by ER stress. We performed an unsupervised k-means clustering (k=15) analysis on the top 5000 genes with the highest variance across different conditions (tunicamycin +/- HDAC inhibitors) (Figure S7b). The clustering captured strong changes resulting from HDAC inhibition (all clusters other than 3 and 13) and tunicamycin treatment (clusters 3 and 13). We first performed GO analysis on tunicamycin-regulated genes and observed a strong increase in ER-stress related pathways (Figure S7d), consistent with existing literature (Liu et al., 2016; Oslowski & Urano, 2011). Strikingly, HDAC inhibition blunted the overall ER stress transcriptional signature and reduced the number of genes induced by tunicamycin alone, providing strong evidence that HDAC inhibition indeed resolves ER stress (Figure S7e-h). This reduction of the ER stress signature could be indirect, reflecting the improved handling of ER stress in neurons treated with HDAC inhibition, or it could be a direct effect of HDAC target genes, or a combination of direct and indirect effects. Interestingly, the expression of different subsets of tunicamycin-induced ER stress genes were shifted in opposite directions upon HDAC inhibition: Cluster 3 was attenuated, and Cluster 13 was enhanced (Figure S7b). To better understand these different groups of tunicamycin-regulated genes, we performed a k-means clustering only on ER stress genes upregulated by tunicamycin (Figure S7i).

This clustering captured three sets of genes that behaved differently upon concurrent HDACi and ER stress. The first cluster was induced by tunicamycin and resolved by HDACi and included genes such as the UPR gene Xbp1 (Figure S7i-j). The second cluster was induced by tunicamycin and enhanced by HDACi and included genes such as the UPR master regulator Eif2ak3/Perk (Figure S7i) and the ERAD gene Derl3 (Figure S7k). The third cluster was induced by tunicamycin and remained elevated in HDACi and included both Manf and Syvn1, both of which we have demonstrated are required for the protective effect in both mouse and *C. elegans* (Figure S7i). Expression of Hspa5, the *hsp-3* ortholog, was also further enhanced in the tunicamycin/HDACi double-treated neurons (Figure 5i), which was consistent with *hsp-3* levels remaining elevated in *ire-1 sin-3* double mutant *C. elegans* neurons (Figure S3d,f).

We next performed PVD-specific RNA-sequencing in *C. elegans* to understand how the loss of *sin-3* or *hda-3* affected transcriptional state.

Previous transcriptomic analyses in PVD have used a non-specific promoter (*ser-2*) that is also expressed in the PDE dopaminergic neurons and the OLL neurons (Taylor et al, 2021). To obtain a pure population of PVD neurons, we developed an intersectional tool to more specifically label only PVD and exclude non-PVD neurons (Figure S8a-b). We used FACS to purify PVD neurons from WT, *sin-3*, and *hda-3* mutant animals and performed RNA-sequencing (Figure S8c-d). We performed GO analysis on the 181 genes that were upregulated by both *hda-3* and *sin-3* (Figure S8e) and found that the most upregulated genes were involved in muscle and gamete development (Figure S8f). These data are consistent with our findings from mouse cortical neurons, in which HDACi resulted in a partial reprogramming of neurons. Together our mouse and *C. elegans* transcriptional data suggested an intriguing hypothesis. In both systems, we observed that HDACi induced upregulation of non-neuronal developmental genes, which were not accompanied by obvious defects in terminal neuronal identity. We hypothesized that this partial reprogramming could represent a transcriptionally “younger” cell state.

To test this hypothesis, we turned to iPSC-derived human neurons. This system was amenable to assessing transcription at multiple timepoints across neuronal differentiation and enabled us to ask whether HDACi could shift neurons backward in developmental time. Human cortical neurons were generated from iPSCs using inducible Neurogenin-2 (NGN2) expression according to standard protocols (Fernandopulle et al, 2018), and RNA was collected at several timepoints across development from Day 0 iPSCs to Day 15 neurons, and from Day 15 neurons treated with TSA (Figure 5d). Successful neuronal differentiation was confirmed by immunostaining (Figure S9a) and transcriptional analysis (Figure S9b-d). To understand how transcriptional state changed during differentiation, we performed principal component analysis which showed a clear time-progression amongst the samples (Figure 5e). PC1 explained 84% of the variance between samples and all developmental time points were clearly separated along the PC1 axis. Neurons treated with TSA at Day 15 were shifted along the PC1 axis to a position equivalent to Day 8 untreated neurons (Figure 5e), and the majority of genes upregulated by TSA were iPSC-enriched (Figure S9f), supporting the hypothesis that TSA reverts neurons to a younger cell state.

We next performed GO analysis (Figure 5f, Figure S9e) and k-means clustering (Figure S9h-i) on genes differentially regulated by TSA in Day 15 neurons. These results were very similar to what was observed in mouse and *C. elegans* neurons: the top GO terms upregulated by TSA were related to development and morphogenesis (Figure 5f), while several GO terms related to neuronal development were downregulated by TSA (Figure S9e). There was strong overlap between TSA-regulated genes in mouse and human neurons (Figure S9g), suggesting that the transcriptional program regulated by HDACi is conserved across species. While proteostasis pathways were not identified by GO analysis, the backup pathway genes involved in ERAD (SYVN1/*sel-11*, SEL1L/*sel-1*), folding (HSPA5/*hsp-3*, MANF/*manf-1*), and UPR (EIF2AK3/*pek-1*, ATF6/*atf-6*) were significantly upregulated by TSA treatment in Day 15 neurons (Figure 5h), suggesting that these are relevant transcriptional targets of HDAC (either direct or indirect). We next asked whether these backup pathway genes represent an important component of the “neuronal rejuvenation” effect. Intriguingly, two of the genes involved in protein folding (MANF and HSPA5) peaked at Day 2 (NPC stage) of neuronal differentiation (Figure 5j and Supplemental Table 4), with expression progressively dropping from Day 5 to Day 15. HDACi treatment returned HSPA5 to Day 2 (Figure 5j) and MANF to Day 5 expression levels (Supplemental Table 4). These data support a model in which HDACi promotes neuronal health by returning neurons to a more flexible, younger state. Neuronal precursors express higher levels of ER folding genes, and the drop in expression over time could leave neurons less equipped to respond to ER stress. HDACi pushes neurons back in developmental time, allowing neurons to express beneficial ER genes that can protect against stress.

## Discussion

Using an unbiased forward genetic screen combined with mechanistic studies in *C. elegans* and mammalian neurons, we uncovered a latent ability of neurons to activate alternative cell biological pathways, which are capable of compensation when preferred growth and membrane trafficking pathways are inactivated. We found that inhibiting class I HDACs improved neuronal health across species and biological contexts by allowing neurons to activate cellular stress-specific resilience pathways. Genetic ablation of *sin-3*/*hda-3* rescued developmental phenotypes in *C. elegans* neurons caused by unfolded protein stress, and pharmacological HDAC inhibition prevented tunicamycin-induced neurodegeneration in mouse cortical neurons. Using genetic and genomic analysis, we identified a network of backup pathway genes that were required for protection, all of which were transcriptionally upregulated by HDACi. Additionally, HDAC inhibition protected *C. elegans* neurons from developmental endosomal trafficking stress, but this protective effect required a distinct set of backup pathway genes. These results demonstrate that neurons usually rely on preferred protein and membrane trafficking pathways, which makes them vulnerable. HDAC inhibition expands the proteostasis repertoires neurons have at their disposal so they can utilize alternative cellular pathways to survive.

HDAC inhibition is beneficial in diverse models of neurodegeneration and neuronal injury (Mai et al., 2009; Shukla & Tekwani, 2020). For example, HDAC inhibitors decreased Aβ plaque number and improved memory impairments in a mouse model of Alzheimer’s disease (Guan et al., 2009), blocked neurodegeneration caused by a mutant polyglutamine repeat containing fragment of the Huntington’s disease protein (McCampbell et al., 2001), and protected against dopaminergic neuron loss in models of Parkinson’s disease (Outeiro et al., 2007). Several potential mechanisms have been proposed, but it is still unresolved how HDAC inhibition provides protection for multiple cell types in different disease and injury conditions. The multitude of transcriptional changes that occur after HDAC inhibition has made it difficult to understand the molecular underpinnings of this protection. We used transcriptomic and loss-of-function analyses to understand both the changes of overall cell state and individual molecular pathways that explain the protective effect. Together our data is consistent with two models: a priming model in which HDACi-treated neurons are poised to allow expression of stress-induced genes, and a rejuvenation model in which neurons have access to earlier developmental programs. The previously described “priming model” proposes that increasing chromatin accessibility via HDACi can prime genomic loci for a robust transcriptional response to an additional stimulus (Graff et al, 2014). Our data are consistent with this model: we find that ER stress alone induces expression of some backup pathway genes, but HDACi enhances expression of ER-stress induced genes required for protection. This suggests that loosened chromatin may provide access to stress-related genes, and that HDACi enables more robust or long-lasting activation of beneficial genes. In addition to the priming model, our data also support a “rejuvenation model,” in which HDACi causes partial reprogramming and pushes neurons back in developmental time. This global upregulation of earlier developmental genes may enable neurons to access additional proteostasis pathways that normally become inaccessible in differentiated and aging neurons.

If HDAC inhibition improves cell survival and functionality and equips neurons to respond at will to distinct sources of stress, why do organisms maintain HDACs in the first place? The expanded functional plasticity afforded by HDAC inhibition may come at a cost. Indeed, *sin-3* mutant animals are sterile, owing to profound germline defects, indicating that HDACs play essential functions during development (Robert et al., 2023), and providing a striking example of antagonistic pleiotropy. Many chromatin modifications have been linked to stress responses and cellular and organismal fitness. For example, activation of the mitochondrial stress response requires the di-methylation of histone H3K9, which silences many genes but also activates protective genes (Tian et al., 2016). Similarly, H3K4me3 level is modulated by reactive oxygen species (ROS) and underlies the benefit of early-life ROS exposure on stress resistance and lifespan (Bazopoulou et al., 2019). Additionally, there are other known examples of pro-degenerative factors that limit the ability of cells to respond to stress. For example, inhibition of the conserved dSarm/Sarm1 molecule dramatically improves axon survival after injury models (Osterloh et al., 2012; Gerdts et al., 2013). Mechanistically, SARM1 triggers axon degeneration locally by NAD+ destruction (Gerdts et al., 2015) and might act as a sensor for metabolic changes that occur after injury, helping decide the fate of the injured axon (Sambashivan & Freeman, 2021). But SARM1 also has an essential function in innate immune signaling and *C. elegans* deficient for the SARM1 ortholog, *tir-1*, are more susceptible to infection (Couillault et al., 2004). Thus, chronic constitutive SARM1 inactivation would be deleterious to an organism whereas acute transient inhibition (perhaps immediately following injury) might be beneficial. Similar principles may apply to HDAC inhibition: while long-term exposure to HDAC inhibitors could result in a detrimental loss of cellular identity, temporary HDAC inactivation during a defined stress could promote beneficial partial rejuvenation.

We have identified a mechanism by which neurons improve cellular flexibility to respond to protein trafficking stressors. We propose that transcriptional flexibility is normally limited during differentiation and development to support cell specialization. Under stressed conditions, this rigid specialization can be detrimental to cells. Increasing transcriptional flexibility improves the ability of cells to activate alternative protein processing pathways and respond to accumulating stress. Cellular chromatin landscapes change during the aging process. A recent study suggests that chromatin flexibility is reduced in aging neurons (Tan et al., 2023), which may contribute to a decreased ability to utilize “backup” pathways to combat cellular stressors. Increasing transcriptional flexibility through chromatin modification may increase brain resilience. It is plausible that additional dormant “backup pathways” exist in neurons and can be activated by epigenetic manipulation to combat diverse neuronal stressors that accumulate with aging, disease, and injury.

## Methods

### *C. elegans* maintenance

*C. elegans* animals were grown on nematode growth media (NGM) plates seeded with OP50 *E. coli*. The wild-type reference strain was N2 Bristol. All experiments were performed at 20°C on L4 animals. Sterile worms were maintained using the hT2 (I;III) or mIn1 (II) balancers and balancer-negative homozygotes were selected for imaging experiments. The *glo-1(zu391) X* background was used for all protein-localization and fluorescent reporter experiments to eliminate autofluorescent gut granules.

The following loss-of-function or hypomorphic alleles were used in this study: LGI: *rab-10(ok1494), dma-1(tm5159), sin-3(wy1123), sin-3(wy1435), sin-3(wy1340), sin-3(wy1336), sin-3(wy1241), sin-3(tm1276), hrdl-1(wy1744), marc-6(wy1747), hda-3(ok1991), eif-2alpha(wy1867);* LGII: *hsp-4(wy2160), hda-2(ok1479), ire-1(wy1569), ire-1(wy1951), hda-5(wy1968), hda-10(tm2996)*; LGIII: *xbp-1(tm2482), der-2(tm6098), rnf-5(tm794), unc-86(wy957), athp-1(wy1748)*; LGIV: *manf-1(wy1746), hlh-30(tm1978), hda-6(tm3436), skn-1(mg570), edem-1(tm5068), mec-3(e1338)*; LGV: *hda-11(tm7128), sel-11(wy1740), sel-1(wy1737), arid-1(wy1749)*; LGX: *hsp-3(wy2124), edem-3(ok1790), atf-6(ok551), pek-1(ok275), hda-4(ok518), glo-1(zu391)*. Deletion alleles were referred to as “Δ” in the figures and text.

The following endogenous fusion proteins or endogenous reporters were used in this study: LGI: *rab-11.1(wy1389[GFP::FLPon(FRT)::rab-11.1]), rab-11.1(wy1947[mScarlet::FLPon(F3)::rab-11.1]), rab-10(wy1616[mScarlet::FLPon(F3)::rab-10]), rab-10(wy1298[GFP::FLPon(FRT)::rab-10]), dma-1(wy1246[DMA-1::GFP]), dma-1(wy1937[DMA-1::GFP]), sin-3(wy1241[sin-3::GFPnovo2]), rab-5(wy1333[GFP::FLPon(FRT)::rab-5]), spcs-1(wy2004[mScarlet::FLPon(F3)::spcs-1]), hda-3(wy1294[hda-3::GFPnovo2])*; LGII: *rab-7(wy1390[GFP::FLPon(FRT)::rab-7]), rab-7(wy1479[mScarlet::FLPon(F3)::rab-7]; LGIV: manf-1(wy2316[P2A::egl-13 NLS::mNG]); LGV: sel-11(wy2271[P2A::egl-13 NLS::mNG]); LGX: hsp-3(wy2314[P2A::egl-13 NLS::mNG]), atf-6(wy2272[P2A::egl-13 NLS::mNG])*.

The following single-copy transgenes were used in this study: LGI: *bmdSi350[pwrt-2::FLP::P2A::H2B::2xmTurq2]*; LGIII: *wySi984[pdes-2::manf-1(cDNA)], wySi987[pdes-2::sel-1(cDNA)], wySi993[pdes-2::rab-11.1(cDNA, S25N)], wySi1013[pdes-2::sel-11(cDNA)], wySi1049[pdes-2::atf-6(cDNA)], wySi1051[pdes-2::pek-1(cDNA)], wySi1047[prab-3::sin-3(cDNA)], wySi1050[peft-3::sin-3(cDNA)], wySi1018[pnhr-81::Cre], wySi1114[pdes-2::hsp-3(cDNA)];* LG V: *wySi937[pdes-2::FLP], wySi960[pdes-2::mcd8-mScarlet], wysi968[pdes-2::mcd8-mScarlet,pdes-2::FLP], wySi996[FLEx-FRT-myr-mScarlet]*, *wySi1008[des-2::cpl-1[W32A,Y35A]:mScarlet]*.

A full list of strains is available in Supplementary Table 1.

### *C. elegans* mutant screen

The *sin-3(wy1123)* and *sin-3(wy1435)* alleles were isolated from an F2 semi-clonal suppressor screen performed in the *rab-10* mutant background. Animals carrying a PVD::GFP transgene and *rab-10* mutation were mutagenized with 50mM ethyl methanesulfonate (EMS). F2 animals were visually screened using a fluorescent compound microscope to identify mutants with restored dendritic branching. Standard SNP-mapping was used to map the location of *wy1123* between +2.3 and +2.88 map units on LGI. Whole genome sequencing was used to identify the causal mutation. The *wy1435* allele failed to complement *wy1123*, and the causal mutation was identified by Sanger sequencing of the *sin-3* locus.

### Construct generation and *C. elegans* genome editing

Endogenous genomic modifications were performed by gonadal microinjection of Cas9 (IDT), gRNA (IDT), tracrRNA (IDT), and PCR-generated or oligonucleotide (ThermoFisher) repair templates. Deletion alleles were generated using two flanking gRNAs and single-stranded oligonucleotide repair templates. Point mutations and small insertions were generated using a single gRNA and single-stranded oligonucleotide repair template. Fluorophore insertions were generated using a single gRNA and PCR product generated from pSK or pSM vectors. All overexpression constructs were created with isothermal assembly (Gibson) and assembled into empty pSK or pSM vector backbones. Single-copy transgene insertions were generated using PCR products generated from these plasmid vectors.

A full list of gRNAs is available in Supplementary Table 1.

### *C. elegans* microscopy

All images were obtained at room temperature in live *C. elegans* animals. L4-stage hermaphrodite animals were anesthetized with 10mM levamisole (Sigma-Aldrich) and mounted on 4% agarose pads. Fluorescently tagged fusion proteins were imaged using a Plan-Apochromat 100x/1.4 NA oil immersion objective on a Zeiss Axio Observer Z1 microscope equipped with a Yokagawa CSU-X1 spinning disk head, a Hamamatsu EM-CCD digital camera, and controlled by MetaMorph (version 7.8.12.0). PVD dendritic morphology was imaged using a C-Apochromat 40x/1.2 NA water immersion objective on a Zeiss Axio Observer Z1 microscope equipped with a Yokogawa CSU-W1 spinning disk unit, a Prime 95B Scientific CMOS camera, and controlled by 3i Slidebook (V6). All image processing and quantification was performed using Fiji. All images were displayed in a standardized orientation (anterior-left, dorsal-top).

To measure dendritic arborization, the full PVD dendrite was imaged in each worm. Z-stacks were acquired at 3-4 positions along each worm. Maximum-intensity projections were generated for each stack and all positions were assembled into one image using the Pairwise Stitching function. Morphology images were straightened and rotated. The Fiji plugin SNT was used to trace the full PVD dendrite and calculate the total length of both primary and higher order dendrites. The length of higher dendrites per 100µm of primary dendrite was calculated for each worm to normalize to primary dendrite length.

To measure the localization and intensity of fluorescently tagged fusion proteins or transcriptional reporters, z-stacks were acquired at the PVD cell soma and proximal primary dendrite. Images were straightened and rotated as z-stacks, and then maximum-intensity and sum-intensity projections were generated. Somatic or nuclear ROIs were drawn in the reference channel (morphology marker). Intensity measurements were obtained from sum-projections, while max-projections were used as display images. Brightness and contrast were adjusted to show relevant features in display images. All images in each experiment were imaged with the same microscope settings and were processed identically for display.

To measure co-localization, an ROI was drawn around the cell soma (for DMA-1/SP12) or the cell body and soma (for DMA-1/RAB-7) and Pearson’s Correlation Coefficient was calculated using the “Coloc 2” Fiji Plugin. All measurements were made on z-stacks, while max-projections were used as display images.

To measure vesicular trafficking, the PVD cell soma and proximal primary dendrite were imaged at a single focal plane. For single color imaging, images were acquired every 200ms, while for two color imaging images were acquired every 600ms. All movies were straightened and rotated for display purposes.

### *C. elegans* statistics and reproducibility

Statistical comparisons were performed in Prism 11 (GraphPad) using one-way ANOVA with Tukey’s correction for multiple comparisons. *C. elegans* data were presented in box-and-whiskers plots that display all points, the median, and the maximum and minimum values. Each data point represents a single worm (one biological replicate). All imaging experiments were repeated across at least three independent sessions.

For PVD morphology measurements, each genotype was assessed individually and the resultant data was subsequently included in multiple experiments for comparison purposes. A summary of instances in which a quantification is included in more than one graph is provided below.

The quantification of wild-type PVD (n=20 individual worms across 5 imaging sessions) that appeared in Figure 1e was repeated in Figure 2b, Figure 3b, Figure S1i, Figure S2g, and Figure S4f.

The quantification of *rab-10(Δ)* PVD (n=20 individual worms across 5 imaging sessions) that appeared in Figure 1e was repeated in Figure 3b, Figure S1c,e, and Figure S4b,f.

The quantification of *rab-10(Δ) sin-3(Δ)* PVD (n=20 individual worms across 5 imaging sessions) that appeared in Figure 1e was repeated in Figure 3b, Figure S1g, and Figure S4b.

The quantification of *ire-1(Δ)* PVD (n=20 individual worms across 6 imaging sessions) that appeared in Figure 1e was repeated in Figure 2b, Figure S1g,i, and Figure S2b,g,k.

The quantification of *ire-1(Δ), sin-3(Δ)* PVD (n=20 individual worms across 6 imaging sessions) that appeared in Figure 1e was repeated in Figure 2b, Figure S1g, Figure S2b,e,k, and Figure S4f.

The quantification of *ire-1(Δ), hda-3(Δ)* PVD (n=20 individual worms across 4 imaging sessions) that appeared in Figure 1e was repeated in Figure S1i.

Quantification of PVD in triple mutants with *ire-1(Δ)* and *sin-3(Δ)* appear in three graphs: individually in Figure 2b, summarized with the full candidate screen in Figure S2b, and as controls for PVD rescue experiments in Figure 2Se. This includes the following genotypes: *ire-1(Δ), sin-3(Δ), sel-11(Δ),* n=20 individual worms across 4 imaging sessions*; ire-1(Δ), sin-3(Δ), sel-1(Δ),* n=20 individual worms across 5 imaging sessions; *ire-1(Δ), sin-3(Δ), manf-1(Δ),* n=20 individual worms across 4 imaging sessions; *ire-1(Δ), sin-3(Δ),atf-6(Δ),* n=20 individual worms across 7 imaging sessions*; ire-1(Δ), sin-3(Δ) pek-1(Δ),* n=20 individual worms across 5 imaging sessions*; ire-1(Δ), sin-3(Δ) hsp-3(cKO),* n=20 individual worms across 5 imaging sessions

For DMA-1::GFP measurements, each genotype was assessed individually and the resultant intensity data was subsequently included in multiple experiments for comparison purposes. The quantifications of DMA-1 intensity in wild-type (n=30 individual worms across 4 imaging sessions) and *ire-1(Δ)* single mutants (n=30 individual worms across 4 imaging sessions) that appeared in Figure 2d were repeated in Figure S2i. A separate set of wild-type controls was used in Figure 3d.

### Mice and primary mouse cortical neurons

Animals were bred and used as approved by the Administrative Panel of Laboratory Animal Care (APLAC) of Stanford University, an institution accredited by the Association for the Assessment and Accreditation of Laboratory Animal Crae (AAALAC). Embryos were harvested from pregnant dams at stage E16.5. Wild-type cultures were generated from C57BL/6 mice.

Primary mouse cortical neurons were dissociated into single-cell suspensions from E16.5 mouse cortices with a papain dissociation system (Worthington Biochemical Corporation). Neurons were seeded onto poly-L-lysine-coated plates (0.1% (wt/vol)) and grown in Neurobasal medium (Gibco) supplemented with B-27 serum-free supplement (Gibco), GlutaMAX, and penicillin-streptomycin (Gibco) in a humidified incubator at 37°C with 5% CO2. Half media changes were performed every 4 or 5 days, or as required. Neurons were plated on 12mm glass coverslips (Carolina Biological Supplies cat#633009) in 24-well plates (300,000 cells/well) and in 12-well plates for RNA extraction (450,000 cells/well). For ER stress induction, tunicamycin (Sigma-Aldrich cat#7765) was added by exchanging the medium for medium containing 1µg/mL tunicamycin, with or without the HDAC inhibitors for 30 hours. The HDAC inhibitors used were 0.1-10 µM Trichostatin A (Sigma-Aldrich cat#T8552) and 0.1-10 µM apicidin (Sigma-Aldrich cat#A8851). Neurites were labeled using antibody against Tuj1/Tubulin β-III (Covance, MRB-435P, 1:1,000 dilution). Alexa-dye-conjugated secondary antibodies (Life Technologies, 1:1,000 dilution) were used.

siRNA oligonucleotide sequences were used to target Syvn1 and Manf (Dharmacon, ON-TARGETplus SMARTpool). For negative controls, a nontarget was used (Dharmacon, ON-TARGETplus Non-Targeting Pool, D-001810). Dissociated cortical neurons were transfected with siRNA 3 days after plating, using the protocol supplied with DharmaFECT 4 (Dharmacon, T-2004-03) and performed as previously described (Maor-Nof et al., 2013) with minor modifications. Briefly, siRNA and the transfection reagent were each diluted separately into NB medium with serum and antibiotics for 5 min; then, the siRNA was added to the medium with the transfection reagent. After an additional 20 min incubation, the transfection reagent siRNA complex was added to the dissociated cells and grown in NB medium without serum and antibiotics. 17 hours later, the transfection reagent was removed by replacing the medium with a complete medium, and the neurons were cultured for an additional 48 hr before treatment with tunicamycin and HDAC inhibitors. The final concentration of the siRNA was 0.1 µM.

### Axonal Quantification *in vitro*

Images of cortical neurons, immuno-stained for β-tubulin, were binarized such that pixels corresponding to axons converted to white while all other regions converted to black. To perform this binarization and differentiate between axons and background in the images, a localized Otsu threshold was used. The Otsu algorithm searches for a threshold that minimizes the variance sum of two or more populations in an image (Otsu, 1979). This gives an exact threshold below which all pixels are considered background. This threshold was then applied to count the number of pixels corresponding to axons in each figure, which serves as the MTs stability index. A punctuated formation of MTs was evident from the cortical explants’ staining; these spots occupy only the higher gray levels in the image and appeared mostly in the Tunicamycin treated, and not in their corresponding untreated neurons. The MT depolymerization index was defined as the ratio of depolymerized axon pixel number to intact axon pixel number. To detect the depolymerized axons, we used an algorithm for counting all the pixels above a certain threshold. To find this threshold, we calculated the probability density function (PDF) of the sum-controlled experiments (untreated neurons), from which the cumulative probability density function (CDF) was extracted. The threshold was set as the value above which there were almost no pixels (less than 0.1%). More than six wells were analyzed for each experimental condition.

### Mouse Statistical Methods

Analyses were performed using R, Microsoft Excel or Prism 8 (GraphPad), and graphs were plotted using Microsoft Excel or Prism 8. For comparisons among different treatment groups, pairwise analyses were conducted by one-way ANOVA. Data represent mean ± SEM unless otherwise noted. Significance is indicated at the figure legends.

### Human iPSC and iPSC-derived neuronal cultures

Human induced pluripotent stem cell (iPSC) research was performed with approval from the Stanford Stem Cell Research Oversight (SCRO) board. Wildtype KOLF2.1J iPSCs (Pantazis et al. 2022) were cultured at 37°C, 5% CO2 in StemFlex media (Thermo Fisher Scientific) on Matrigel-coated wells, and passaged 1:20 every 3-4 days with ReLeSR (STEMCELL Technologies).

To generate a doxycycline-inducible Neurogenin-2 (NGN2) cell line for cortical neuron differentiation, PiggyBac-mediated genomic insertion of PB-TO-oNGN2 (Addgene #198397) into KOLF2.1J iPSCs was performed by co-transfection of PB-TO-oNGN2 and EF1a-transposase using Lipofectamine Stem Transfection Reagent (Thermo Fisher Scientific, Cookson et al. 2022 from *iNDI protocols.io*). iPSCs were treated with 2 µg/mL puromycin for 2 weeks, to select for a stable polyclonal cell line expressing PB-TO-oNGN2, with >97% cells BFP-positive for the NGN2 construct by flow cytometry analysis. The KOLF2.1J PB-TO-oNGN2 line tested negative for mycoplasma contamination, had no karyotype abnormalities by optical genome mapping, and displayed normal iPSC morphology.

Cortical neurons were generated using doxycycline-inducible expression of NGN2 (Fernandopulle et al. 2018, Lara Flores 2023 from *iNDI protocols.io*). On Day 0, iPSCs were dissociated with Accutase (Innovative Cell Technologies) and plated at a density of 500K cells per each 6-well on Matrigel-coated wells, in Induction Media (DMEM/F-12 HEPES with 1x GlutaMAX, 1x NEAA, 1x N2, and 2 µg/mL doxycycline) with 1x CEPT (Chroman 1, Emricasan, Polyamine supplement, Trans-ISRIB). On Days 1 and 2, media was exchanged for fresh Induction Media (without CEPT).

On Day 3, NPCs were dissociated with Accutase and plated into Maturation Media (50% DMEM/F-12 HEPES : 50% BrainPhys, with 1x N21-MAX, 10 ng/mL BDNF, GDNF, and NT-3, 1 µM uridine and FUDR, 1 µg/mL mouse laminin, and 2 µg/mL doxycycline) at a density of 150K cells per each well in a 24-well format. Prior to plating, wells were coated with Poly-D-Lysine (0.1 mg/mL, Thermo Fisher Scientific) and mouse laminin (1 mg/mL, SouthernBiotech), including acid-etched glass coverslips (Carolina Biologicals) for microscopy or no coverslips (for RNAseq experiments). Half-volume media changes with Maturation Media were performed on Days 4, 7, 10, and 14. From Day 10 onwards, Maturation Media contained 100% BrainPhys (STEMCELL technologies).

TSA drug treatment (10 µM, 7 hr) or DMSO control treatment was performed on Day 15. Samples were either collected for RNAseq (described below) or fixed for immunocytochemistry experiments. Neurons were fixed by gently adding an equal volume of 4% paraformaldehyde (Electron Microscopy Services) in PBS to the cell culture media (final concentration 2%) for 5 min, followed by gentle aspiration and additional fixation in 4% PFA in PBS for 10 min at room temperature. Neurons were washed in PBS, permeabilized in 0.1% Triton X-100 in PBS for 5 min, and blocked in 10% goat serum in PBS for 1 hr. Primary antibody staining was performed overnight at 4°C, with Map2 (PhosphoSolutions, 1:2000), Tuj1 (BioLegend, 1:1000), and Acetyl-Histone H2B Lys5 (Cell Signaling, 1:400). Neurons were washed in PBS and incubated in secondary antibodies (Alexa Fluor goat anti-chicken 647, anti-mouse 568, and anti-rabbit 488) for 2 hr. Neurons were washed in PBS, counterstained with Hoechst 33342, and mounted on glass slides in Aqua/Poly-Mount. Coverslips were imaged using a 63x/1.2 NA water immersion objective on a Zeiss Axio Observer Z1 microscope equipped with a Yokogawa CSU-W1 spinning disk unit, a Prime 95B Scientific CMOS camera, 405 nm / 488 nm / 561 nm / and 640 nm lasers, and controlled by 3i Slidebook.

### Mouse RNA-sequencing

RNA samples were collected from 4 experimental replicates (batches of cultures). Primary mouse neurons were treated for 7 hours with either tunicamycin, tunicamycin and HDAC inhibitors, or left untreated. For RNA extraction and isolation, neurons were lysed in RLT buffer from the QIAGEN RNeasy Plus Micro Kit (QIAGEN), according to the manufacturer’s protocol. In brief, cells were lysed, homogenized, and loaded on a genomic DNA (gDNA) eliminator column. After removal of genomic DNA, RNA was purified using RNeasy spin columns. RNA quantity and purity were determined by optical density measurements of OD260/280 and OD230/260 using a NanoDrop spectrophotometer. Structural integrity of the total RNA was assessed with the Agilent 2100 Bioanalyzer using the Agilent Total RNA Nano Chip assay (Agilent Technologies). mRNA libraries were prepared for Illumina paired-end sequencing with an Agilent SureSelect Strand Specific RNA-Seq Library Preparation Kit on an Agilent Bravo Automated Liquid Handling Platform. Libraries were sequenced on an Illumina NextSeq sequencer at the Chan Zuckerberg Initiative (CZI).

### Human RNA sequencing

RNA samples were collected from 3 experimental replicates (independent rounds of neuron induction) with 7 conditions in each experiment (Day 0, Day 2, Day 5, Day 8, Day 11, Day 15 + DMSO, Day 15 + TSA 7 hr 10 µM). RNA was extracted using the QIAGEN RNeasy Plus Micro Kit (QIAGEN).

Library preparation and RNA sequencing was performed by Novogene. RNA was measured with a NanoDrop spectrophotometer and Agilent Bioanalyzer 2100, to confirm RNA quantity and quality. mRNA was purified from total RNA using poly-T oligo-attached magnetic beads, and libraries were prepared for sequencing using the unstranded NEBNext Ultra RNA Library Prep Kit. RNA sequencing was performed using an Illumina platform to generate paired-end 150 bp reads.

### *C. elegans* RNA sequencing

*C. elegans* were synchronized and cultured at 20°C, and animals were harvested as Day 1 adults. Two biological replicates from independent plates were harvested for each genotype, which were each split into two technical replicates (sorted and processed into libraries independently) for a total of four technical replicates. *C. elegans* were dissociated to isolate nuclei as previously described (Gao et al., 2024) and fluorescent positive nuclei were sorted using a Sony SH800 cell sorter. cDNA was synthesized from sorted nuclei using the Smart-SCRB protocol (Dong et al., 2024), a modified SCRB-seq approach employing template-switching reverse transcription with Maxima H Minus reverse transcriptase and KAPA HiFi PCR amplification. Sequencing libraries were prepared using the Nextera XT DNA Library Preparation Kit (Illumina) and sequenced on an Illumina NextSeq 2000 to generate paired-end reads.

### RNA-seq data analysis

Mouse and human FASTQ files were processed with fastp 1.0.1 (Chen et al., 2018) to remove adapters, poly-N sequences, and low-quality sequences. Reads were aligned to the human genome (GRCh38.115) or mouse genome (GRCm39.114) using STAR 2.7.10b (Dobin et al., 2013) and reads per gene were counted using HTSeq 2.0.1 (Putri et al., 2022). *C. elegans* FASTQ files were processed with fastp (Chen et al., 2018) to remove adapters, poly-N sequences, and short reads (<15 bp). Read quality was assessed with FastQC and MultiQC. Reads were aligned to the *C. elegans* genome (WormBase WS287) using STAR (Dobin et al., 2013), and gene-level read counts were generated with featureCounts (Liao et al., 2014) in paired-end mode. One wild-type replicate (MG4) was excluded from downstream analysis due to a low mapping rate (<1%), likely caused by RNA degradation.

DESeq2 1.48.1 (Love et al., 2014) was used to obtain normalized read counts and perform differential expression analysis. Differentially expressed genes were defined with abs(log2FoldChange > 1) and p-adjusted < 0.05. Principal component analysis (PCA) was performed on the top 500 most variable genes in the dataset.

Biological process GO enrichment analysis was performed on lists of differentially expressed genes, using the Gene Ontology Resource (Gene Ontology Consortium, 2025), using all pass-filtered genes as background. GO terms with fold-enrichment > 1.5 were sorted by p-value and bubble plots of the top GO terms were generated (Tang et al., 2023).

Gene expression profiles were visualized using heatmaps, with expression values standardized via row-wise Z-score scaling and partitioned into distinct groups through K-means clustering.

Differential gene expression of ER stress genes was visualized with volcano plots, with abs log2fc > 1 and p-adjusted < 0.05 as the cutoffs. The list of 315 ER stress genes was defined by genes associated with any of the 16 relevant GO terms (ERAD pathway, response to unfolded protein, response to endoplasmic reticulum stress, and similar).

Stacked bar graphs were generated by comparing human TSA vs DMSO differentially expressed genes (abs log2fc > 1 and p-adjusted < 0.05) to a second dataset (Day 15 vs Day 0, or mouse TSA vs DMSO), using a lower threshold for the second dataset (abs log2fc > 0.5 and p-adjusted < 0.05).

## Data Availability

Raw RNAseq data will be made available on the Gene Expression Omnibus. For convenient analysis, tables of raw reads, normalized reads, and differential gene expression are included in Supplemental Tables 2 (mouse), 3 (*C. elegans*), and 4 (human). Human iNeuron timecourse data visualization is available at https://kang-shen-lab.shinyapps.io/ineuron_timecourse/.

## Supplemental Movie legends

**Supplemental Movie 1. DMA-1 localizes to RAB-10 vesicles in wild type PVD.**

Representative movie of co-labeled DMA-1::GFP and mScarlet::RAB-10 trafficking in PVD neuron. Two-color images acquired every 600ms.

**Supplemental Movie 2. DMA-1 is mis-trafficked to RAB-7 late endosomes.**

Representative movies of co-labeled DMA-1::GFP and mScarlet::RAB-7 trafficking in wild type, *rab-10* single mutant, and *rab-10, sin-3* double mutant PVD neurons. Two-color images acquired every 600ms.

**Supplemental Movie 3. DMA-1 is mis-trafficked to RAB-11.1 recycling endosomes.**

Representative movies of co-labeled DMA-1::GFP and mScarlet::RAB-11.1 trafficking in wild type and *rab-10* single mutant PVD neurons. Two-color images acquired every 600ms.

**Supplemental Movie 4. RAB-11.1 trafficking is altered by loss of *rab-10* and *sin-3*.**

Representative movies of GFP::RAB-11.1 trafficking in wild type, *sin-3* single mutant, and *rab-10, sin-3* double mutant PVD neurons. Images acquired every 200ms.

**Supplemental Movie 5. RAB-10 is trafficked to higher order dendrites.**

Representative movie of GFP::RAB-10 trafficking in wild type PVD neuron. Images acquired every 200ms.

## Acknowledgments

We thank colleagues in the Shen laboratory for helpful discussion and Victor Paat for lab support. For worm strains, we thank Kelsie Eichel, the National Bioresource Project for the nematode, the International C. elegans Gene Knockout Consortium, and the CGC, which is funded by NIH Office of Research Infrastructure Programs (P40 OD010440). PB-TO-oNGN2 was a gift from iPSC Neurodegenerative Disease Initiative (iNDI) & Michael Ward (Addgene plasmid #198397). We thank Mark Edgley for assistance in sequencing balanced mutants. We thank the Howard Hughes Medical Institute at Janelia Research Campus Quantitative Genomics Core Facility (RRID:SCR_022694) and Lihua Wang for reverse transcription, library construction, and sequencing of *C. elegans* samples. We thank Youchen Guan and Lang Ding from the Meng Wang lab for assistance with worm husbandry and nuclei isolation. This work is supported by The Phil and Penny Knight Initiative for Brain Resilience at Stanford University. W.A.H. is supported by NIH NRSA Fellowship F32 NS129942. K.S is an investigator of the Howard Hughes Medical Institute. A.D.G. is supported by NIH (grants R35NS097263, U54NS123743, R01AG064690) and Target ALS. A.D.G. is a Chan Zuckerberg Biohub – San Francisco Investigator.

## Author contributions

C.A.T. conceptualization, methodology, *C. elegans* experiments, data analysis, data visualization, writing—original draft

M.M.N. mouse experiments, data analysis, writing—review and editing

W.A.H. human iPSC neuron experiments, RNAseq data analysis for human, mouse, and worm, writing—review and editing

S.M.G *C. elegans* sorting and sequencing experiments, data analysis

X.L. human iPSC and neuron cultures

E.M.R. RNAseq data analysis for mouse

A.H. *C. elegans* experiments

F.Z. and B.S. iPSC resources: generation of KOLF2.1J PB-TO-oNGN2 iPSC line

C.Y. *C. elegans* resources

M.C.W. supervision

A.D.G. supervision, writing—review and editing

K.S. methodology, supervision, writing—original draft

---

**Supplemental Figure 1.**
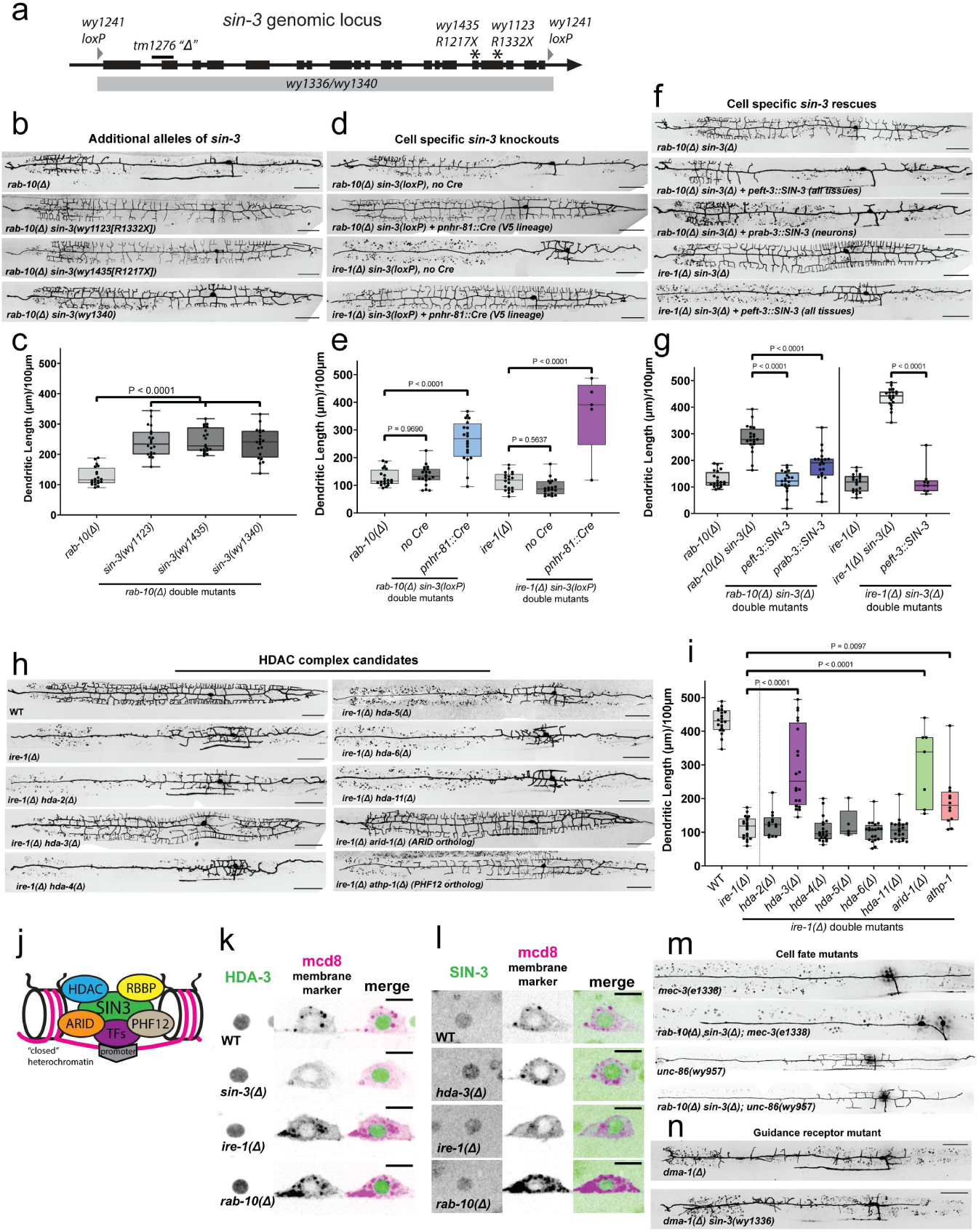
SIN-3 functions cell-autonomously in a nuclear complex with HDA-3. Related to Figure 1. (a) Schematic of the sin-3 genomic locus depicting all alleles used in this study. (b) Representative images of PVD morphology in *rab-10* single mutant and double mutants of *rab-10* and three *sin-3* alleles, *wy1123, wy1435,* and *wy1340*. (c) Quantification of total dendritic length per 100 µm of primary dendrite for genotypes shown in b. (d) Representative images of PVD morphology in double mutants of *rab-10* or *ire-1* and floxed *sin-3* with and without Cre. (e) Quantification of total dendritic length per 100 µm of primary dendrite for genotypes shown in d. (f) Representative images of PVD morphology in double mutants of *rab-10 sin-3* or *ire-1 sin-3* with single-copy overexpression of SIN-3(cDNA). (g) Quantification of total dendritic length per 100 µm of primary dendrite for genotypes shown in f. (h) Representative images of PVD morphology in wild type, *ire-1* single mutants, and double mutants of *ire-1* and candidate HDAC complex genes. All PVD morphology scale bars in a, d, f, and h are 50 µm. (i) Quantification of total dendritic length per 100 µm of primary dendrite for genotypes shown in h. (j) Schematic of predicted *C. elegans* SIN-3/HDAC complex. (k) Representative images of endogenous HDA-3::GFP with a red membrane marker (*pdes-2::mcd8::mScarlet*) used to visualize PVD morphology in wild-type, *sin-3, ire-1* and *rab-10* mutants. Scale bars are 5 µm. (k) Representative images of endogenous SIN-3::GFP with a red membrane marker (*pdes-2::mcd8::mScarlet*) used to visualize PVD morphology in wild-type, *sin-3, ire-1* and *rab-10* mutants. Scale bars are 5 µm. (m) Representative images of PVD morphology in cell fate mutants *mec-3* or *unc-86*, and triple mutants of *rab-10 sin-3* with *mec-3* or *unc-86*. (n) Representative images of PVD morphology in guidance receptor mutant *dma-1* or double mutant of *dma-1 sin-3*. PVD morphology scale bars are 50 µm. All statistical comparisons were performed using one-way ANOVA with Tukey’s correction for multiple comparisons.

**Supplemental Figure 2.**
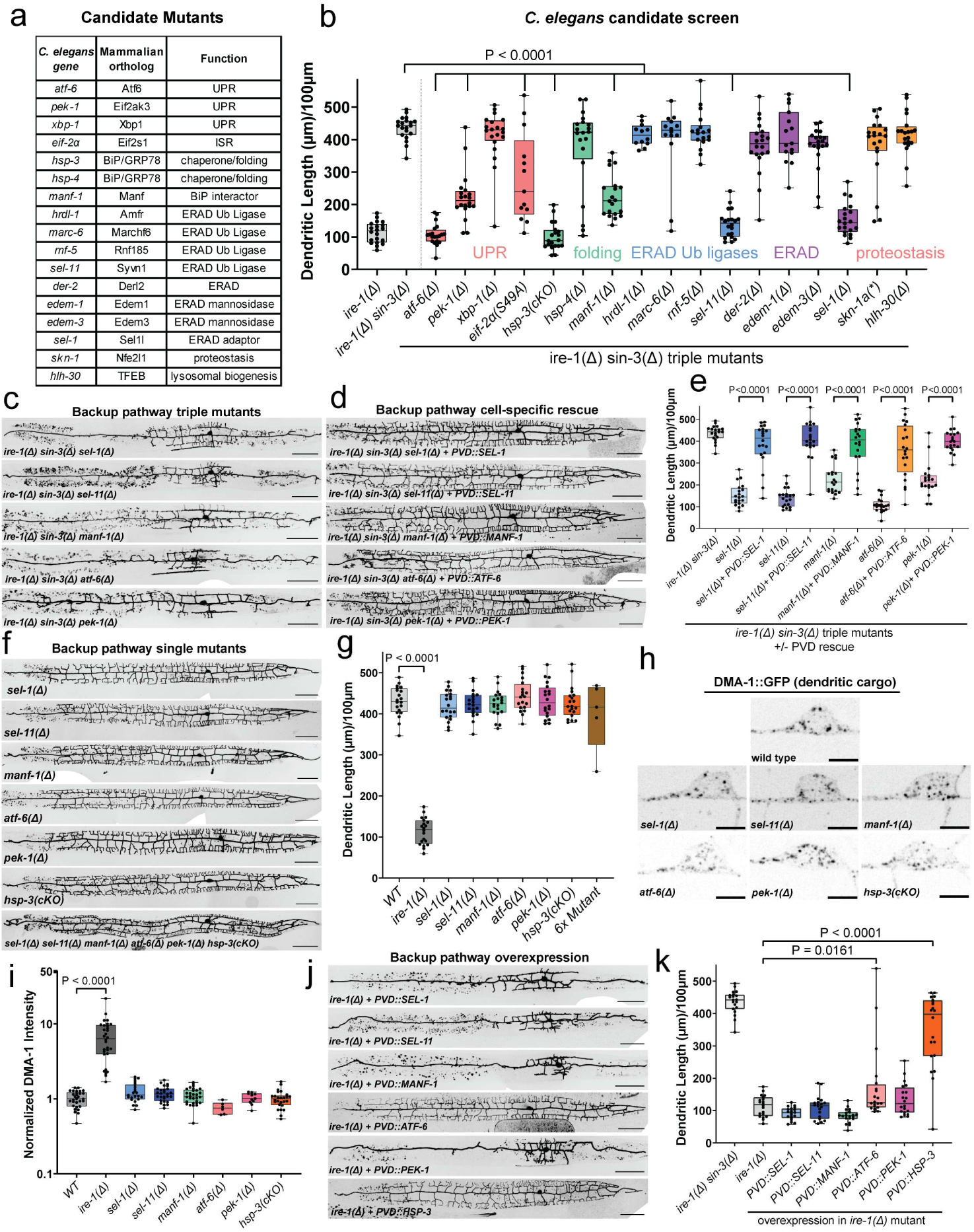
Candidate screen reveals backup pathway genes, none of which are required for normal development. Related to Figure 2. (a) List of genes tested in candidate screen. (b) Quantification of total dendritic length per 100 µm of primary dendrite for triple mutants of *ire-1 sin-3* with all genes listed in a. (c) Representative images of PVD morphology in triple mutants of *ire-1 sin-3* with *sel-1, sel-11, manf-1, atf-6, pek-1,* or *hsp-3.* Scale bars are 50 µm. (d) Representative images of PVD morphology in triple mutants of *ire-1 sin-3* with *sel-1, sel-11, manf-1, atf-6, pek-1,* or *hsp-3* with cell-specific rescue constructs of the corresponding cDNA (i.e. PVD::SEL-1 expressed in *ire-1 sin-3 sel-1* mutant). Scale bars are 50 µm. (e) Quantification of total dendritic length per 100 µm of primary dendrite for all genotypes in c-d. (f) Representative images of PVD morphology in single mutants of *sel-1, sel-11, manf-1, atf-6, pek-1, hsp-3,* or a sextuple mutant of all above genes. Scale bars are 50 µm. (g) Quantification of total dendritic length per 100 µm of primary dendrite for all genotypes in f. (h) Representative images of endogenous DMA-1 GFP in soma of wild-type, *sel-1, sel-11, manf-1, atf-6, pek-1,* or *hsp-3* single mutants. Scale bars are 5 µm. (i) Quantification of DMA-1::GFP intensity for all genotypes in h. All genotypes were normalized to the median of wild-type. (j) Representative images of PVD morphology in *ire-1* single mutants with cell-specific expression of individual backup pathway genes. Scale bars are 50 µm. (k) Quantification of total dendritic length per 100 µm of primary dendrite for all genotypes in j. All statistical comparisons were performed using one-way ANOVA with Tukey’s correction for multiple comparisons.

**Supplemental Figure 3.**
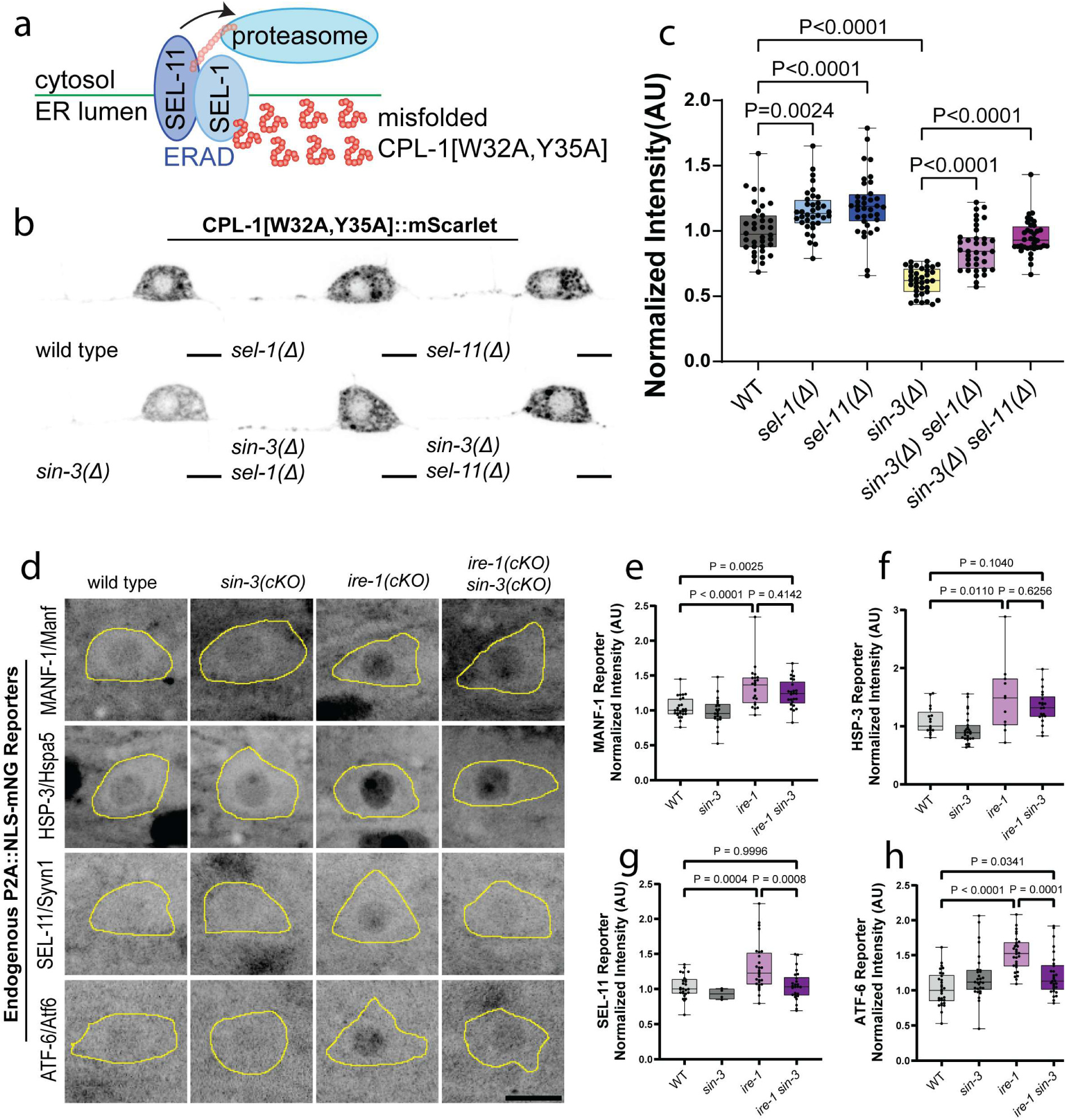
Loss of *sin-3* boosts ERAD activity and alters *ire-1* transcriptional response. Related to Figure 2. (a) Schematic of ERAD sensor CPL-1[W32A,Y35A], a misfolded protein whose clearance is dependent upon ERAD. (b) Representative images of ERAD substrate CPL-1[W32A,Y35A]::mScarlet in wild-type, *sin-3*, *sel-1,* or *sel-11* single mutants, and double mutants of *sin-3 sel-1* or *sin-3 sel-11*. Scale bars are 5 µm. (c) Quantification of CPL-1[W32A,Y35A]::mScarlet intensity for all genotypes in b. All genotypes were normalized to the median of wild-type. (d) Representative images of endogenous transcriptional reporters of MANF-1, HSP-3, SEL-11, or ATF-6 in wild-type, *sin-3* single mutant, *ire-1* single mutant, or *ire-1 sin-3* single mutants. PVD soma is outlined in yellow. Scale bar is 5 µm and applies to all images in grid. (e) Quantification of MANF-1 reporter intensity for all genotypes shown in d. (f) Quantification of HSP-3 reporter intensity for all genotypes shown in d. (g) Quantification of SEL-11 reporter intensity for all genotypes shown in d. (h) Quantification of ATF-6 reporter intensity for all genotypes shown in d. All statistical comparisons were performed using one-way ANOVA with Tukey’s correction for multiple comparisons.

**Supplemental Figure 4.**
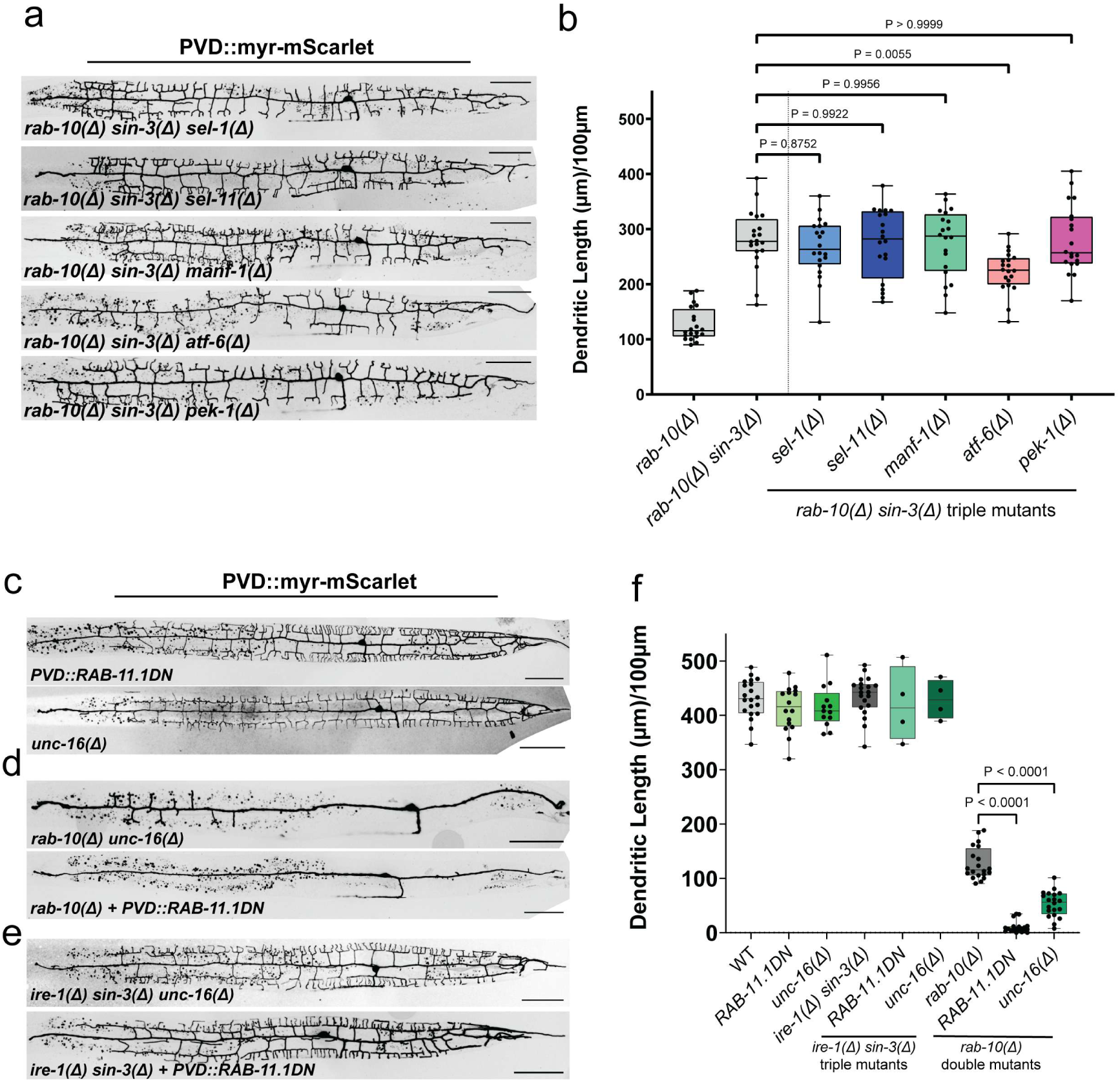
Distinct backup pathways are required to protect against different developmental stressors. Related to Figure 3. (a) Representative images of PVD morphology in triple mutants of *rab-10 sin-3* with *sel-1, sel-11, manf-1, atf-6,* or *pek-1.* Scale bars are 50 µm. (b) Quantification of total dendritic length per 100 µm of primary dendrite for all genotypes in a. (c) Representative images of PVD morphology in single mutants of RAB-11.1(DN) or *unc-16*. (d) Representative images of PVD morphology in double mutants of *rab-10* with RAB-11.1(DN) or *unc-16*. (e) Representative images of PVD morphology in triple mutants of *ire-1 sin-3* with RAB-11.1(DN) or *unc-16*. All PVD morphology scale bars are 50 µm. (f) Quantification of total dendritic length per 100 µm of primary dendrite for all genotypes in c-e. All statistical comparisons were performed using one-way ANOVA with Tukey’s correction for multiple comparisons.

**Supplemental Figure 5.**
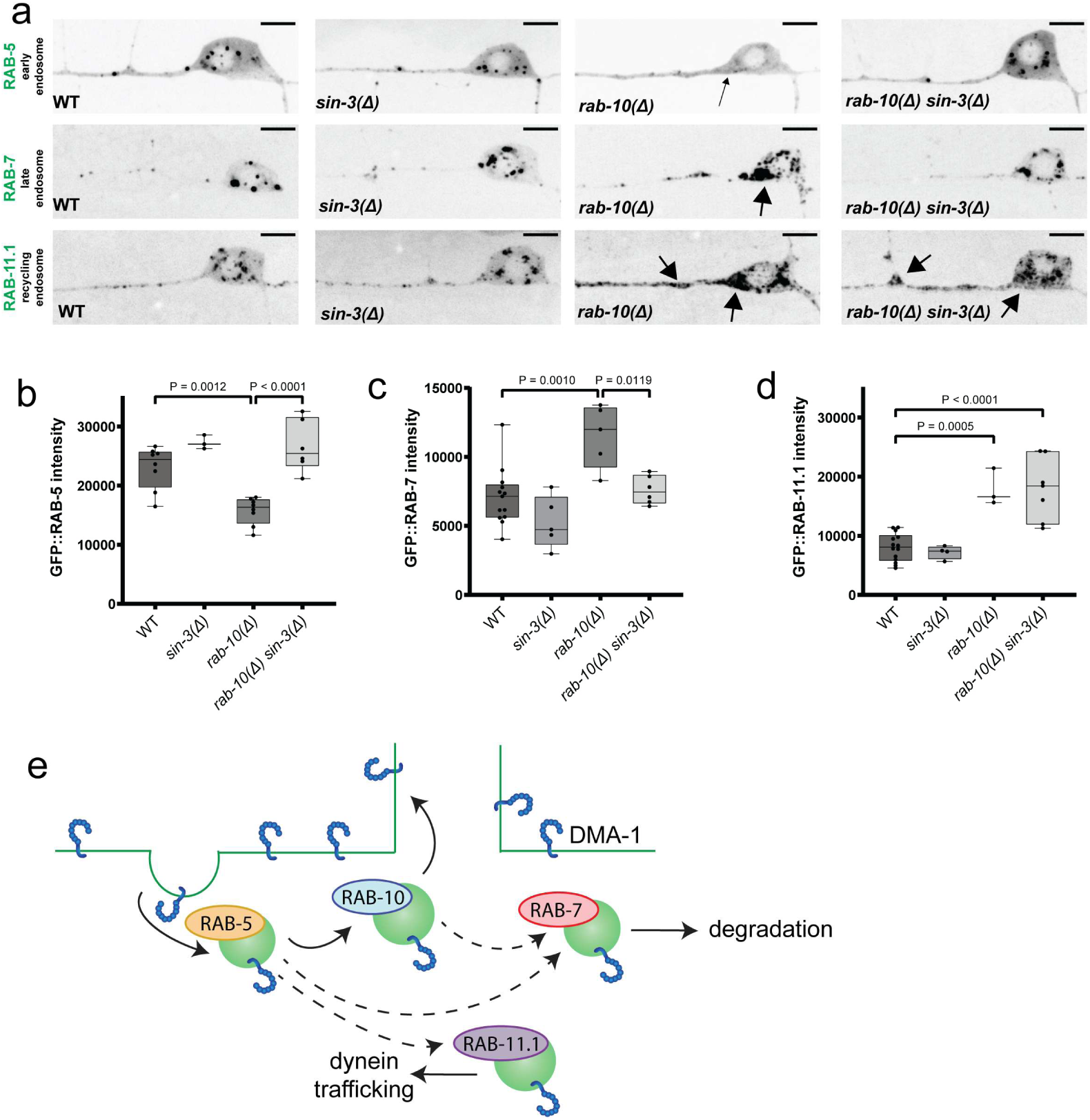
Distribution of Rab endosomes is altered by *rab-10* and *sin-3.* Related to Figure 3. (a) Representative images of endogenous GFP::RAB-5 (top), endogenous GFP::RAB-7 (middle), and endogenous GFP::RAB-11.1 (bottom) in wild-type, *sin-3* and *rab-10* mutants, and *rab-10, sin-3* double mutants. Scale bars are 5 µm. (b) Quantification of GFP::RAB-5 intensity in PVD soma and dendrite for genotypes in a. (c) Quantification of GFP::RAB-7 intensity in the PVD soma and dendrite for genotypes in a. (d) Quantification of GFP::RAB-11.1 intensity in the PVD soma and dendrite for genotypes in a. (e) Schematic of routes of endosomal traffic in PVD. All statistical comparisons were performed using one-way ANOVA with Tukey’s correction for multiple comparisons.

**Supplemental Figure 6.**
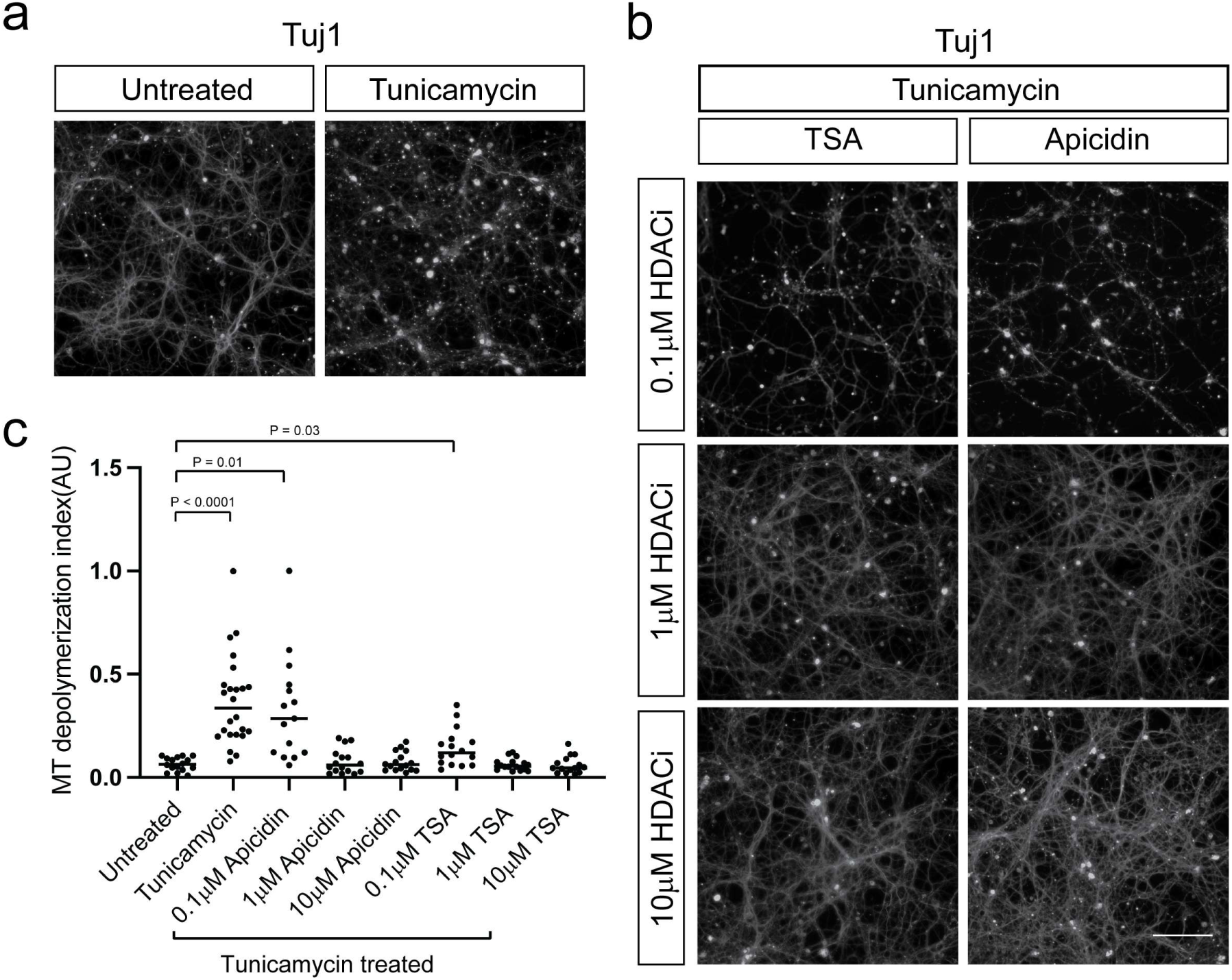
HDACi acts in a dose-dependent manner. Related to Figure 4. (a) Representative images of tubulin βΙΙΙ staining in mouse cortical neurons treated with or without tunicamycin. (b) Representative images of tubulin βΙΙΙ staining in mouse cortical neurons co-treated with tunicamycin and either apicidin or TSA. (c) Quantification of microtubule depolymerization index determined from tubulin βΙΙΙ staining. All statistical comparisons were performed using one-way ANOVA with Tukey’s correction for multiple comparisons. Scale bar is 100 µm.

**Supplemental Figure 7.**
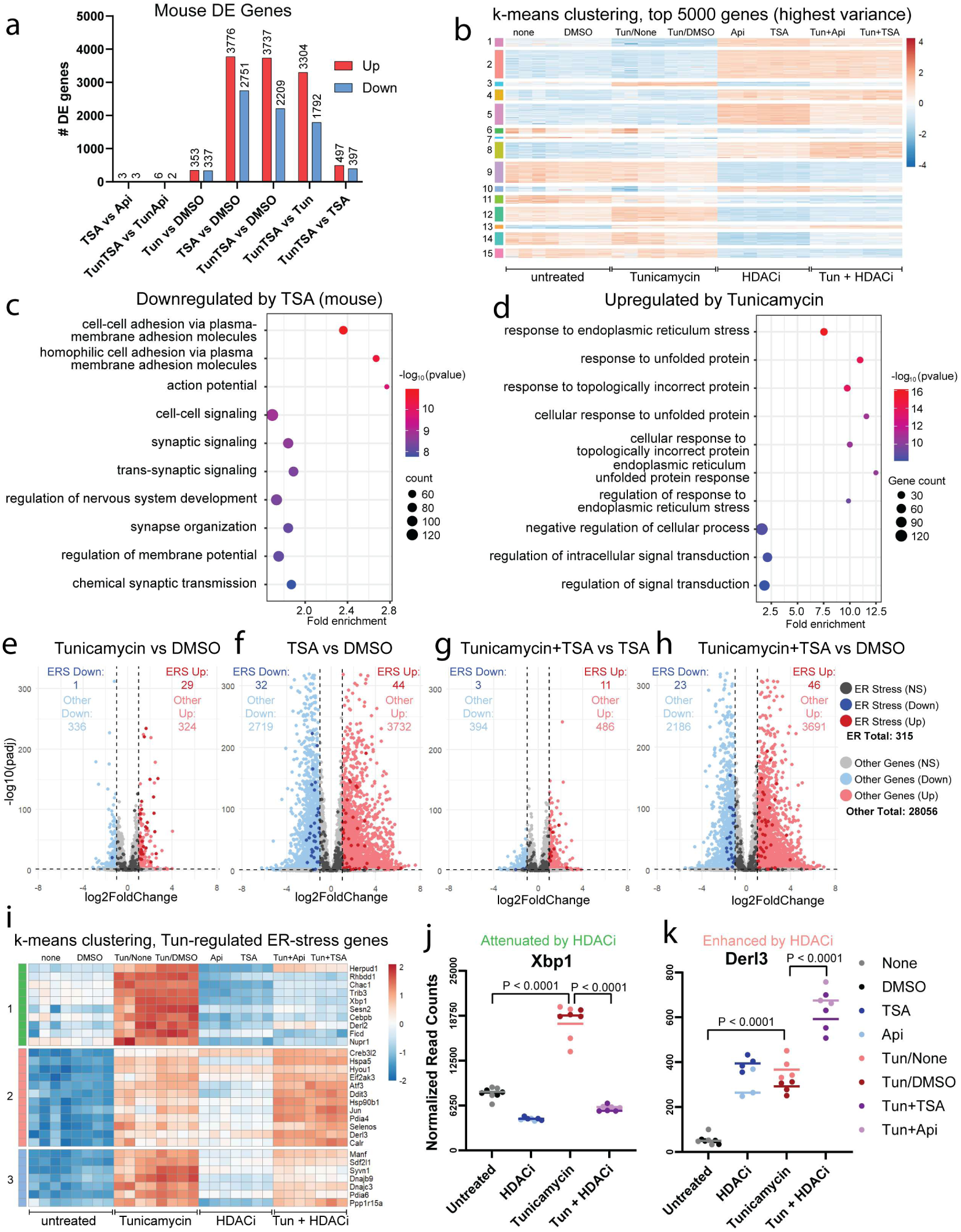
HDACi modulates the tunicamycin-induced transcriptional landscape in mouse cortical neurons. Related to Figure 5. (a) Total number of differentially expressed genes between different conditions. (b) k-means clustering performed on the top 5000 genes of highest variance, k=15. (c) GO analysis of genes downregulated by TSA. (d) GO analysis of genes upregulated by tunicamycin. (e-h) Volcano plots showing up- and down-regulated genes in Tunicamycin vs. DMSO (e), TSA vs. DMSO (f), tunicamycin + TSA vs. TSA (g), and tunicamycin + TSA vs. DMSO (h). (i) k-means clustering performed on ER stress genes upregulated by tunicamycin. (j) Normalized gene counts for select ER stress gene Xbp1, which is induced by tunicamycin and attenuated by HDACi. (k) Normalized gene counts for select ERAD gene Derl3, which is induced by tunicamycin and enhanced by HDACi.

**Supplemental Figure 8.**
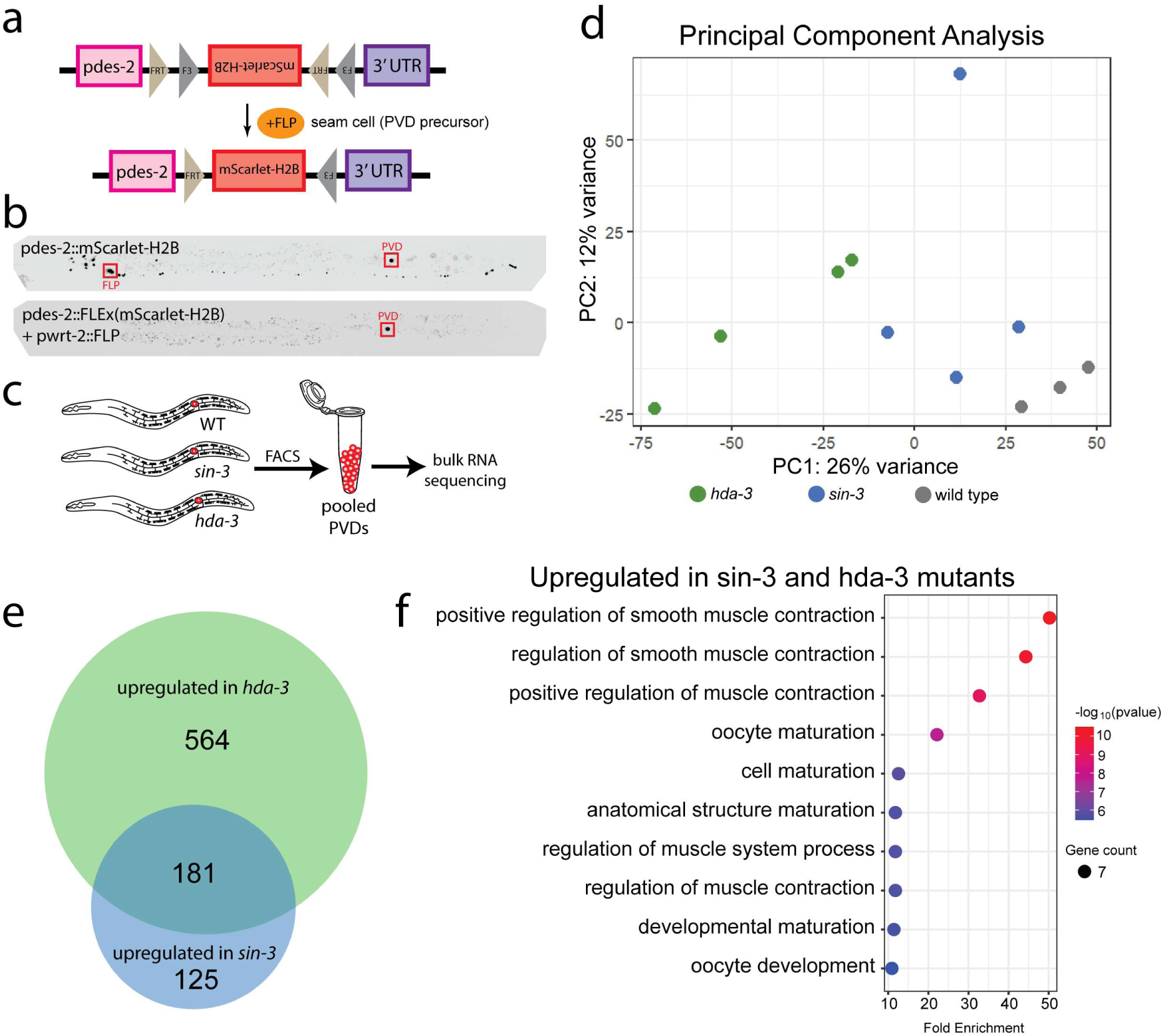
Loss of *sin-3/hda-3* modifies transcriptional cell state in *C. elegans.* Related to Figure 5. (a) Schematic of intersectional FLEx/FLP design to accomplish cell-specific PVD nuclear labeling. (b) Representative images of nuclei labeled by *des-2* (top) or the intersectional FLEx/FLP driven by the seam cell promoter *pwrt-2* (bottom). (c) Schematic of RNA-sequencing experiment in *C. elegans*. (d) Principal component analysis. (e) Total number of upregulated genes in *hda-3* and *sin-3* mutants. (f) GO analysis of genes upregulated in both *hda-3* and *sin-3* mutants.

**Supplemental Figure 9.**
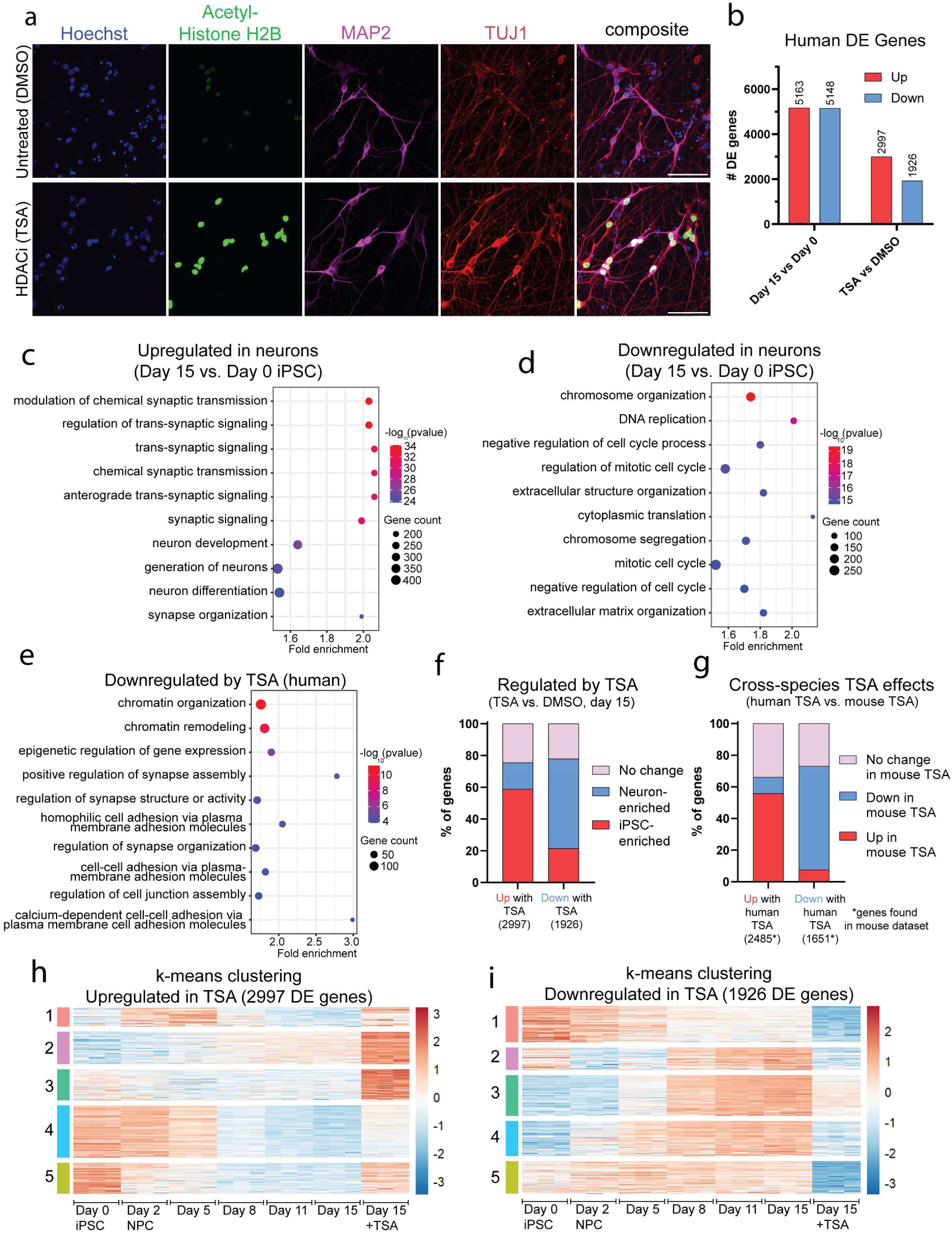
HDACi regulates a similar transcriptional program in human and mouse neurons. Related to Figure 5. (a) Representative images of iPSC-derived neurons with or without HDACi, stained for nuclei (Hoescht), acetyl-histone H2B (Lys5), somatodendritic compartments (MAP2), and all neurites (TUJ1). (b) Total number of differentially expressed genes in Day 15 neurons compared to Day 0 iPSCs or to TSA treated neurons. (c) GO analysis of genes upregulated in Day 15 neurons, as compared to Day 0 iPSCs. (d) GO analysis of genes downregulated in Day 15 neurons, as compared to Day 0 iPSCs. (e) GO analysis of genes downregulated by TSA in human neurons. (f) Proportion of neuron-enriched and iPSC-enriched genes regulated by TSA. (g) Comparison of TSA-regulated genes across mouse and human neurons. Only genes with orthologs found in the mouse dataset were included in the analysis. (h) k-means clustering of 2997 genes upregulated by TSA, k=5. (i) k-means clustering of 1926 genes downregulated by TSA, k=5.

## References

1. Balch, W. E., Morimoto, R. I., Dillin, A., & Kelly, J. W. (2008). Adapting proteostasis for disease intervention. *Science (New York*, N.Y.), 319(5865), 916–919. 10.1126/science.1141448

2. Bazopoulou, D., Knoefler, D., Zheng, Y., Ulrich, K., Oleson, B. J., Xie, L., Kim, M., Kaufmann, A., Lee, Y.-T., Dou, Y., Chen, Y., Quan, S., & Jakob, U. (2019). Developmental ROS individualizes organismal stress resistance and lifespan. Nature, 576(7786), 301–305. 10.1038/s41586-019-1814-y

3. Bradner, J. E., West, N., Grachan, M. L., Greenberg, E. F., Haggarty, S. J., Warnow, T., & Mazitschek, R. (2010). Chemical phylogenetics of histone deacetylases. Nature Chemical Biology, 6(3), 238–243. 10.1038/nchembio.313

4. Chen, S., Zhou, Y., Chen, Y., & Gu, J. (2018). fastp: an ultra-fast all-in-one FASTQ preprocessor. *Bioinformatics (Oxford*, England), 34(17), i884–i890. 10.1093/bioinformatics/bty560

5. Choong, C.-J., Sasaki, T., Hayakawa, H., Yasuda, T., Baba, K., Hirata, Y., Uesato, S., & Mochizuki, H. (2016). A novel histone deacetylase 1 and 2 isoform-specific inhibitor alleviates experimental Parkinson’s disease. Neurobiology of Aging, 37, 103–116. 10.1016/j.neurobiolaging.2015.10.001

6. Cookson, M., Ward, M., & Flores, E.L. (2023). iNDI PiggyBac-TO-hNGN2 transfection protocol Version 1. protocols.io. 10.17504/protocols.io.rm7vzxb22gx1/v1

7. Couillault, C., Pujol, N., Reboul, J., Sabatier, L., Guichou, J.-F., Kohara, Y., & Ewbank, J. J. (2004). TLR-independent control of innate immunity in Caenorhabditis elegans by the TIR domain adaptor protein TIR-1, an ortholog of human SARM. Nature Immunology, 5(5), 488–494. 10.1038/ni1060

8. Dobin, A., Davis, C. A., Schlesinger, F., Drenkow, J., Zaleski, C., Jha, S., Batut, P., Chaisson, M., & Gingeras, T. R. (2013). STAR: ultrafast universal RNA-seq aligner. *Bioinformatics (Oxford*, England), 29(1), 15–21. 10.1093/bioinformatics/bts635

9. Dong, P., Zhang, S., Gandin, V., Xie, L., Wang, L., Lemire, A. L., Li, W., Otsuna, H., Kawase, T., Lander, A. D., Chang, H. Y., & Liu, Z. J. (2024). Cohesin prevents cross-domain gene coactivation. Nature genetics, 56(8), 1654–1664. 10.1038/s41588-024-01852-1

10. Eichel, K., Uenaka, T., Belapurkar, V., Lu, R., Cheng, S., Pak, J. S., Taylor, C. A., Südhof, T. C., Malenka, R., Wernig, M., Özkan, E., Perrais, D., & Shen, K. (2022). Endocytosis in the axon initial segment maintains neuronal polarity. Nature, 609(7925), 128–135. 10.1038/s41586-022-05074-5

11. Fernandopulle, M. S., Prestil, R., Grunseich, C., Wang, C., Gan, L., & Ward, M. E. (2018). Transcription Factor-Mediated Differentiation of Human iPSCs into Neurons. Current protocols in cell biology, 79(1), e51. 10.1002/cpcb.51

12. Finger, A., Knop, M., & Wolf, D. H. (1993). Analysis of two mutated vacuolar proteins reveals a degradation pathway in the endoplasmic reticulum or a related compartment of yeast. European journal of biochemistry, 218(2), 565–574. 10.1111/j.1432-1033.1993.tb18410.x

13. Flores, E.L., Qi, A., Reilly, L., Santiana, M., Ward, M., & Cookson, M. iNDI Transcription Factor-NGN2 differentiation of human iPSC into cortical neurons Version 2. protocols.io. 10.17504/protocols.io.n2bvj3owblk5/v1

14. Fu, M. M., & Holzbaur, E. L. (2014). Integrated regulation of motor-driven organelle transport by scaffolding proteins. Trends in cell biology, 24(10), 564–574. 10.1016/j.tcb.2014.05.002

15. Gao, S. M., Qi, Y., Zhang, Q., Guan, Y., Lee, Y. T., Ding, L., Wang, L., Mohammed, A. S., Li, H., Fu, Y., & Wang, M. C. (2024). Aging atlas reveals cell-type-specific effects of pro-longevity strategies. Nature aging, 4(7), 998–1013. 10.1038/s43587-024-00631-1

16. Gerdts, J., Brace, E. J., Sasaki, Y., DiAntonio, A., & Milbrandt, J. (2015). SARM1 activation triggers axon degeneration locally via NAD+ destruction. *Science (New York*, N.Y.), 348(6233), 453–457. 10.1126/science.1258366

17. Gerdts, J., Summers, D. W., Sasaki, Y., DiAntonio, A., & Milbrandt, J. (2013). Sarm1-mediated axon degeneration requires both SAM and TIR interactions. The Journal of Neuroscience: The Official Journal of the Society for Neuroscience, 33(33), 13569–13580. 10.1523/JNEUROSCI.1197-13.2013

18. Gräff, J., Joseph, N. F., Horn, M. E., Samiei, A., Meng, J., Seo, J., Rei, D., Bero, A. W., Phan, T. X., Wagner, F., Holson, E., Xu, J., Sun, J., Neve, R. L., Mach, R. H., Haggarty, S. J., & Tsai, L. H. (2014). Epigenetic priming of memory updating during reconsolidation to attenuate remote fear memories. Cell, 156(1-2), 261–276. 10.1016/j.cell.2013.12.020

19. Guan, J.-S., Haggarty, S. J., Giacometti, E., Dannenberg, J.-H., Joseph, N., Gao, J., Nieland, T. J. F., Zhou, Y., Wang, X., Mazitschek, R., Bradner, J. E., DePinho, R. A., Jaenisch, R., & Tsai, L.-H. (2009). HDAC2 negatively regulates memory formation and synaptic plasticity. Nature, 459(7243), 55–60. 10.1038/nature07925

20. Heiman, M. G., & Bülow, H. E. (2024). Dendrite morphogenesis in Caenorhabditis elegans. Genetics, 227(2), iyae056. 10.1093/genetics/iyae056

21. Hetz, C., & Saxena, S. (2017). ER stress and the unfolded protein response in neurodegeneration. Nature Reviews. Neurology, 13(8), 477–491. 10.1038/nrneurol.2017.99

22. Hoozemans, J. J. M., van Haastert, E. S., Nijholt, D. A. T., Rozemuller, A. J. M., Eikelenboom, P., & Scheper, W. (2009). The unfolded protein response is activated in pretangle neurons in Alzheimer’s disease hippocampus. The American Journal of Pathology, 174(4), 1241–1251. 10.2353/ajpath.2009.080814

23. Hubbert, C., Guardiola, A., Shao, R., Kawaguchi, Y., Ito, A., Nixon, A., Yoshida, M., Wang, X.-F., & Yao, T.-P. (2002). HDAC6 is a microtubule-associated deacetylase. Nature, 417(6887), 455–458. 10.1038/417455a

24. Janczura, K. J., Volmar, C.-H., Sartor, G. C., Rao, S. J., Ricciardi, N. R., Lambert, G., Brothers, S. P., & Wahlestedt, C. (2018). Inhibition of HDAC3 reverses Alzheimer’s disease-related pathologies in vitro and in the 3xTg-AD mouse model. Proceedings of the National Academy of Sciences of the United States of America, 115(47), E11148–E11157. 10.1073/pnas.1805436115

25. Kurtishi, A., Rosen, B., Patil, K. S., Alves, G. W., & Møller, S. G. (2019). Cellular Proteostasis in Neurodegeneration. Molecular Neurobiology, 56(5), 3676–3689. 10.1007/s12035-018-1334-z

26. Liao, Y., Smyth, G. K., & Shi, W. (2014). featureCounts: an efficient general purpose program for assigning sequence reads to genomic features. Bioinformatics (Oxford, England), 30(7), 923–930. 10.1093/bioinformatics/btt656

27. Lippincott-Schwartz, J., Roberts, T. H., & Hirschberg, K. (2000). Secretory protein trafficking and organelle dynamics in living cells. Annual Review of Cell and Developmental Biology, 16, 557–589. 10.1146/annurev.cellbio.16.1.557

28. Liu, C.-L., Zhong, W., He, Y.-Y., Li, X., Li, S., & He, K.-L. (2016). Genome-wide analysis of tunicamycin-induced endoplasmic reticulum stress response and the protective effect of endoplasmic reticulum inhibitors in neonatal rat cardiomyocytes. Molecular and Cellular Biochemistry, 413(1–2), 57–67. 10.1007/s11010-015-2639-0

29. Liu, O. W., & Shen, K. (2011). The transmembrane LRR protein DMA-1 promotes dendrite branching and growth in C. elegans. Nature Neuroscience, 15(1), 57–63. 10.1038/nn.2978

30. Love, M. I., Huber, W., & Anders, S. (2014). Moderated estimation of fold change and dispersion for RNA-seq data with DESeq2. Genome biology, 15(12), 550. 10.1186/s13059-014-0550-8

31. Mai, A., Rotili, D., Valente, S., & Kazantsev, A. G. (2009). Histone deacetylase inhibitors and neurodegenerative disorders: Holding the promise. Current Pharmaceutical Design, 15(34), 3940–3957. 10.2174/138161209789649349

32. Maor-Nof, M., Homma, N., Raanan, C., Nof, A., Hirokawa, N., & Yaron, A. (2013). Axonal pruning is actively regulated by the microtubule-destabilizing protein kinesin superfamily protein 2A. Cell Reports, 3(4), 971–977. 10.1016/j.celrep.2013.03.005

33. McCampbell, A., Taye, A. A., Whitty, L., Penney, E., Steffan, J. S., & Fischbeck, K. H. (2001). Histone deacetylase inhibitors reduce polyglutamine toxicity. Proceedings of the National Academy of Sciences of the United States of America, 98(26), 15179–15184. 10.1073/pnas.261400698

34. Nijholt, D. A. T., van Haastert, E. S., Rozemuller, A. J. M., Scheper, W., & Hoozemans, J. J. M. (2012). The unfolded protein response is associated with early tau pathology in the hippocampus of tauopathies. The Journal of Pathology, 226(5), 693–702. 10.1002/path.3969

35. Oslowski, C. M., & Urano, F. (2011). Measuring ER stress and the unfolded protein response using mammalian tissue culture system. Methods in Enzymology, 490, 71–92. 10.1016/B978-0-12-385114-7.00004-0

36. Osterloh, J. M., Yang, J., Rooney, T. M., Fox, A. N., Adalbert, R., Powell, E. H., Sheehan, A. E., Avery, M. A., Hackett, R., Logan, M. A., MacDonald, J. M., Ziegenfuss, J. S., Milde, S., Hou, Y.-J., Nathan, C., Ding, A., Brown, R. H., Conforti, L., Coleman, M., … Freeman, M. R. (2012). dSarm/Sarm1 is required for activation of an injury-induced axon death pathway. Science (New York, N.Y.), 337(6093), 481–484. 10.1126/science.1223899

37. Otsu, N. (1979) A Threshold Selection Method from Gray-Level Histograms. IEEE Transactions on Systems, Man, and Cybernetics, 9, 62–66. 10.1109/TSMC.1979.4310076

38. Outeiro, T. F., Kontopoulos, E., Altmann, S. M., Kufareva, I., Strathearn, K. E., Amore, A. M., Volk, C. B., Maxwell, M. M., Rochet, J.-C., McLean, P. J., Young, A. B., Abagyan, R., Feany, M. B., Hyman, B. T., & Kazantsev, A. G. (2007). Sirtuin 2 inhibitors rescue alpha-synuclein-mediated toxicity in models of Parkinson’s disease. Science (New York, N.Y.), 317(5837), 516–519. 10.1126/science.1143780

39. Pantazis, C. B., Yang, A., Lara, E., McDonough, J. A., Blauwendraat, C., Peng, L., Oguro, H., Kanaujiya, J., Zou, J., Sebesta, D., Pratt, G., Cross, E., Blockwick, J., Buxton, P., Kinner-Bibeau, L., Medura, C., Tompkins, C., Hughes, S., Santiana, M., Faghri, F., … Merkle, F. T. (2022). A reference human induced pluripotent stem cell line for large-scale collaborative studies. Cell stem cell, 29(12), 1685–1702.e22. 10.1016/j.stem.2022.11.004

40. Putri, G. H., Anders, S., Pyl, P. T., Pimanda, J. E., & Zanini, F. (2022). Analysing high-throughput sequencing data in Python with HTSeq 2.0. *Bioinformatics (Oxford*, England), 38(10), 2943–2945. 10.1093/bioinformatics/btac166

41. Rivieccio, M. A., Brochier, C., Willis, D. E., Walker, B. A., D’Annibale, M. A., McLaughlin, K., Siddiq, A., Kozikowski, A. P., Jaffrey, S. R., Twiss, J. L., Ratan, R. R., & Langley, B. (2009). HDAC6 is a target for protection and regeneration following injury in the nervous system. Proceedings of the National Academy of Sciences of the United States of America, 106(46), 19599–19604. 10.1073/pnas.0907935106

42. Robert, V. J., Caron, M., Gely, L., Adrait, A., Pakulska, V., Couté, Y., Chevalier, M., Riedel, C. G., Bedet, C., & Palladino, F. (2023). SIN-3 acts in distinct complexes to regulate the germline transcriptional program in Caenorhabditis elegans. *Development (Cambridge*, England), 150(21), dev201755. 10.1242/dev.201755

43. Salzberg, Y., Coleman, A. J., Celestrin, K., Cohen-Berkman, M., Biederer, T., Henis-Korenblit, S., & Bülow, H. E. (2017). Reduced Insulin/Insulin-Like Growth Factor Receptor Signaling Mitigates Defective Dendrite Morphogenesis in Mutants of the ER Stress Sensor IRE-1. PLoS Genetics, 13(1), e1006579. 10.1371/journal.pgen.1006579

44. Sambashivan, S., & Freeman, M. R. (2021). SARM1 signaling mechanisms in the injured nervous system. Current Opinion in Neurobiology, 69, 247–255. 10.1016/j.conb.2021.05.004

45. Seto, E., & Yoshida, M. (2014). Erasers of histone acetylation: The histone deacetylase enzymes. Cold Spring Harbor Perspectives in Biology, 6(4), a018713. 10.1101/cshperspect.a018713

46. Shi, R., Ho, X. Y., Tao, L., Taylor, C. A., Zhao, T., Zou, W., Lizzappi, M., Eichel, K., & Shen, K. (2024). Stochastic growth and selective stabilization generate stereotyped dendritic arbors. bioRxiv: The Preprint Server for Biology, 2024.05.08.591205. 10.1101/2024.05.08.591205

47. Shukla, S., & Tekwani, B. L. (2020). Histone Deacetylases Inhibitors in Neurodegenerative Diseases, Neuroprotection and Neuronal Differentiation. Frontiers in Pharmacology, 11, 537. 10.3389/fphar.2020.00537

48. Singh, R., Kaur, N., Choubey, V., Dhingra, N., & Kaur, T. (2024). Endoplasmic reticulum stress and its role in various neurodegenerative diseases. Brain Research, 1826, 148742. 10.1016/j.brainres.2023.148742

49. Smith, C. J., Watson, J. D., Spencer, W. C., O’Brien, T., Cha, B., Albeg, A., Treinin, M., & Miller, D. M. (2010). Time-lapse imaging and cell-specific expression profiling reveal dynamic branching and molecular determinants of a multi-dendritic nociceptor in C. elegans. Developmental Biology, 345(1), 18–33. 10.1016/j.ydbio.2010.05.502

50. Suelves, N., Kirkham-McCarthy, L., Lahue, R. S., & Ginés, S. (2017). A selective inhibitor of histone deacetylase 3 prevents cognitive deficits and suppresses striatal CAG repeat expansions in Huntington’s disease mice. Scientific Reports, 7(1), 6082. 10.1038/s41598-017-05125-2

51. Tan, L., Shi, J., Moghadami, S., Parasar, B., Wright, C. P., Seo, Y., Vallejo, K., Cobos, I., Duncan, L., Chen, R., & Deisseroth, K. (2023). Lifelong restructuring of 3D genome architecture in cerebellar granule cells. *Science (New York*, N.Y.), 381(6662), 1112–1119. 10.1126/science.adh3253

52. Tang, D., Chen, M., Huang, X., Zhang, G., Zeng, L., Zhang, G., Wu, S., & Wang, Y. (2023). SRplot: A free online platform for data visualization and graphing. PloS one, 18(11), e0294236. 10.1371/journal.pone.0294236

53. Taylor, C. A., Yan, J., Howell, A. S., Dong, X., & Shen, K. (2015). RAB-10 Regulates Dendritic Branching by Balancing Dendritic Transport. PLoS Genetics, 11(12), e1005695. 10.1371/journal.pgen.1005695

54. Taylor, S. R., Santpere, G., Weinreb, A., Barrett, A., Reilly, M. B., Xu, C., Varol, E., Oikonomou, P., Glenwinkel, L., McWhirter, R., Poff, A., Basavaraju, M., Rafi, I., Yemini, E., Cook, S. J., Abrams, A., Vidal, B., Cros, C., Tavazoie, S., Sestan, N., … Miller, D. M., 3rd (2021). Molecular topography of an entire nervous system. Cell, 184(16), 4329–4347.e23. 10.1016/j.cell.2021.06.023

55. Tian, Y., Garcia, G., Bian, Q., Steffen, K. K., Joe, L., Wolff, S., Meyer, B. J., & Dillin, A. (2016). Mitochondrial Stress Induces Chromatin Reorganization to Promote Longevity and UPR(mt). Cell, 165(5), 1197–1208. 10.1016/j.cell.2016.04.011

56. Tsalik, E. L., Niacaris, T., Wenick, A. S., Pau, K., Avery, L., & Hobert, O. (2003). LIM homeobox gene-dependent expression of biogenic amine receptors in restricted regions of the C. elegans nervous system. Developmental Biology, 263(1), 81–102. 10.1016/s0012-1606(03)00447-0

57. Way, J. C., & Chalfie, M. (1989). The mec-3 gene of Caenorhabditis elegans requires its own product for maintained expression and is expressed in three neuronal cell types. Genes & Development, 3(12A), 1823–1833. 10.1101/gad.3.12a.1823

58. Wei, X., Howell, A. S., Dong, X., Taylor, C. A., Cooper, R. C., Zhang, J., Zou, W., Sherwood, D. R., & Shen, K. (2015). The unfolded protein response is required for dendrite morphogenesis. eLife, 4, e06963. 10.7554/eLife.06963

59. Wiseman, R. L., Mesgarzadeh, J. S., & Hendershot, L. M. (2022). Reshaping endoplasmic reticulum quality control through the unfolded protein response. Molecular Cell, 82(8), 1477–1491. 10.1016/j.molcel.2022.03.025

60. Zou, W., Yadav, S., DeVault, L., Nung Jan, Y., & Sherwood, D. R. (2015). RAB-10-Dependent Membrane Transport Is Required for Dendrite Arborization. PLoS Genetics, 11(9), e1005484. 10.1371/journal.pgen.1005484

